# Second generation of LNP based mRNA vaccine leads to a T cell–inflamed tumor microenvironment favorable for improving PD-1/PD-L1 blocking therapy and long-term immunity in a cold tumor model

**DOI:** 10.1101/2024.07.24.604918

**Authors:** Carole Fournier, Marion Mercey-Ressejac, Valentin Derangère, Amal Al Kadi, David Rageot, Christine Charrat, Alexis Leroy, Marie Escudé, Séverine Escaich, François Ghiringhelli, Thomas Decaens, Fabrice P. Navarro, Evelyne Jouvin-Marche, Patrice N. Marche

**Affiliations:** Institute for Advanced Biosciences, Univ. Grenoble Alpes, Inserm U 1209, CNRS UMR 5309, Grenoble, France; Univ. Grenoble Alpes, CEA, LETI, Technologies for Healthcare and Biology Division, Microtechnologies for Living Systems Interactions Research Unit, F-38000 Grenoble, France; INSERM U1231, Equipe TIRECS, 21000 Dijon France; Université de Bourgogne, 21000 Dijon France; Centre de Lutte contre le Cancer Georges François Leclerc, Plateforme de Transfert en Biologie du Cancer, 21000 Dijon

## Abstract

The delivery of mRNA-based cancer vaccines has demonstrated significant promise in triggering antitumor immune responses. With the aim of using them in combination with other immunotherapies already used in the clinical appropriately, the modifications of the intratumoral immune microenvironment needs to be deeply characterized. We have shown that the second generation of lipid nanoparticles (LNPs), nanostructured lipid carriers (so-called Lipidots^®^), are able to vector protein antigens and nucleic acids. Here, we test Lipidots^®^ for the delivery of mRNA encoding OVA antigen and eliciting a specific antitumor immune response. We demonstrate *in vitro* that our LNPs deliver mRNA into dendritic cells (DCs), when complexed with mRNA, activate DCs through the TLR4/8 and ROS signaling pathways and induce specific CD4^+^ and CD8^+^ T cell activation. Our vaccinal strategy exhibits significant antitumor efficacy both in the context of tumor prevention and as a therapeutic vaccine in B16OVA and E.G7-OVA cold tumors. The LNP-Ova mRNA vaccine induces a profound intratumoral remodeling of the innate and adaptive immunity associated with an increase in the gene expression of chemokines (*Cxcl10, Cxcl11, Cxcl9*) involved in CD8^+^ T cell attraction. Additionally, the vaccine induces the establishment of an escape mechanism mediated by PD-1/PDL-1 axis, making it an adjuvant therapy for optimized responses to the blocking of this signaling pathway. Finally, the combination of vaccine and anti-PD-1 therapy achieves a much higher rate of complete responses and memory immune responses compared to monotherapies. Our work demonstrates the capability of Lipidots^®^ as an effective platform for the development of preventive and therapeutic vaccines against cancer based on mRNA delivery and that combination with other immunotherapies such as immune checkpoint blockers could counter tumor resistance and promote long-term antitumor immunity.

## Introduction

The combination of immunosuppression in the tumor microenvironment (TME) and low immunogenicity of cancer cells is an important feature that hinders effective immune-mediated tumor regression. Tumors explore a variety of mechanisms to evade immune detection including expression of immune inhibitory ligands such as programmed death ligand 1 (PD-L1), repression of tumor antigen presentation and release of immunosuppressive molecules such as TGF-β (1). To rearm an effective antitumoral immune response, immune checkpoint blockade (ICB), which notably targets the key signaling axis involving programmed death-1 (PD-1) and its ligands PD-L1 and PD-L2, is the most commonly used immunotherapy in cancer patients and demonstrates remarkable beneficial effects in various solid malignancies (2–5). Despite its clinical success, many patients do not respond optimally to ICB due to primary or acquired resistance (6). Complementary therapeutic strategies that aim to make the tumor hotter by stimulating T-effector-cell infiltration are needed to maximize the clinical effectiveness of ICB (7–9).

The impairment of naïve T cell priming caused by the activation of cancer cell-specific oncogenic pathways, which leads to ineffective intratumoral recruitment of dendritic cells (DCs), is associated with ICB resistance in melanoma and hepatocellular cancer models (10,11). In weakly immunogenic and/or immune desert tumors, therapeutic cancer vaccine by eliciting a specific T cell response that promotes intratumoral T-cell trafficking and because of its low toxicity would be a leading treatment to be combined with ICB (12–14). With the approval of COVID-19 mRNA vaccines using first generation of lipid nanoparticle (LNP) delivery platforms by US FDA, the safety and efficacy of this therapeutic technology were broadly demonstrated (15,16). Importantly, mRNA-based cancer vaccines have several advantages including their good tolerability, non-infectious composition, in addition to their fast and low-cost production (17). Among the diverse nanocarrier pharmaceutical systems, lipid-based nanodelivery systems have become the most common used vectors for mRNA cancer vaccine because of high mRNA preservation and cellular uptake (18). Following administration, LNP-mRNA based vaccine is mainly internalized by antigen presenting cells (APCs), including DCs, which translate the mRNA-encoding tumor-associated antigen while undergoing maturation and activation that benefits the subsequent generation of the adaptive immunity (19,20). Although this therapy has not yet been approved for standard treatment in oncology, multiple clinical trials of mRNA vaccines are currently on going (13) and encouraging results have already been obtained for melanoma and pancreatic cancer (21,22). Several preclinical studies have demonstrated the antitumor efficacy of vaccines delivering mRNA coding for OVA or tumor antigens (23–33), however, they have rarely explored the reshaping of the tumor immune microenvironment, focusing more on evaluating the peripheral response, or the anticancer memory immunity. Since these two aspects are essential for understanding the mechanisms of sensitivity and resistance to immunotherapies, we particularly addressed them.

We have developed cationic nanostructured lipid carriers, named Lipidots^®^, formulated with biocompatible and FDA-approved ingredients (wax, oil, lecithin, PEG-stearate) that have been extensively characterized for physicochemical aspects (34–38). This LNP, so called Lipidot^®^, has a natural tropism toward lymph nodes (39) and are capture by DCs from mesenteric lymph nodes following an intraperitoneal injection (36). Moreover, their capacity to deliver into bacteria, fibroblasts, DCs and macrophages nucleic acids, which are negatively charged and allow electrostatic interactions with Lipidots^®^, has been evidence *in vitro* (37,40,41). Based on that, we investigated whether LNP-mRNA coding ovalbumin (OVA) complex can be transfected in primary murine DCs and trigger an OVA specific T cell response. We also evaluated its capacity to induce a specific antitumoral immune response in murine melanoma and lymphoma models expressing OVA (B16OVA and E.G7-OVA) and to improve the effectiveness of anti-PD-1 blocking antibody in B16OVA tumors which are poorly infiltrated with immune cells and resistant to anti-PD-1 therapy. First, we show that our mRNA-based vaccine strategy activates DCs through Toll-Like Receptor (TLR) and reactive oxygen species (ROS) signaling *in vitro* and is efficiently translated and processed to induce OVA-specific CD8^+^ and CD4^+^ T cell activations. Subsequently, we demonstrate that LNP-Ova mRNA-based vaccine decreases tumor growth in association with the promotion of a T cell–inflamed tumor microenvironment, characterized by active IFN-γ signaling, cytotoxic effector molecules and antigen presentation that are all essential features for anti-PD-1 responsiveness (8,42). Combining mRNA-based vaccine with anti-PD-1 partially prevents the intratumoral accumulation of CD8^+^ T cells co-expressing PD-1 and TIM-3, increases the early production of granzyme B by intratumoral CD4^+^ T cells, improves survival and leads to a higher rate of total tumor regression. Finally, mRNA-based vaccine +/- anti-PD-1-treated mice surviving the first B16OVA tumor engraftment are totally resistant to tumor re-challenge expressing OVA (B16OVA) while partially resistant when tumor cells do not express OVA (B16F10) implying the development of immunologic memory and antigen spreading.

## Materials and methods

### Mice

Wild type C57BL/6 mice were purchased from Charles River laboratories (Saint-Germain sur l’Arbresle, France). OT-I and OT-II mice were bred at «Plateforme de Haute Technologie Animale (PHTA)» University of Grenoble-Alpes core facility (Grenoble, France). Female mice greater than 9 weeks of age were matched by age and randomly attributed to the different treatment groups. The number of mice in each treated group for each experiment is indicated in the legend of the figures. No mice were excluded based on preestablished criteria, and randomization was applied only immediately pretreatment to ensure similar mean tumor size at the start of therapy experiments. Animal use and care were approved by the Animal Experiments Ethics Committee “Comité d’Ethique pour l’Expérimentation Animale no.#12, Cometh-Grenoble” and approved by the French Ministry of Research (#32695-2021081311583272 v3).

### Cell culture

Mouse B16OVA (melanoma) and E.G7-OVA (lymphoma) cell lines were maintained in humidified air with 5% CO_2_ at 37°C in the appropriate culture media. Both cell lines were grown in Roswell Park Memorial Institute (RPMI) 1640 w/L-glutamine supplemented with 10% (vol/vol) heat-inactivated fetal bovine serum (FBS,), 1 mM Sodium Pyruvate (Gibco) and 1% Penicillin Streptomycin (P/S). B16OVA media was completed with 1% Non-Essential Amino Acids (NEAA) and E.G7-OVA media with 10 mM HEPES, 0.05 mM 2-mercaptoethanol and 0.4 mg/ml G418 (Gibco).

For bone marrow derived DCs (BMDCs), bone marrow was isolated from the tibias and femurs of female C57BL/6 mice (6 weeks old) and processed into a single cell suspension as described in a previous study (43). GR1 positive cells and erythrocytes were removed following incubation with Ly-6G/Ly-6C (BD Pharmingen, #553125) and TER-119 (BD Pharmingen, #553672) antibodies, and the remaining negatively sorted cells were isolated using Dynabeads isolation kit (ThermoFisher, #11047) by magnetic cell sorting. Then, the sorted cells were cultured at 5 × 10^5^ cells/ml into a 100-mm standard tissue culture dish (Falcon™ #353003) with 15 mL of complete Iscove’s modified Dulbecco’s medium (IMDM containing 10% heat-inactivated FBS, 1% NEAA, 1% sodium pyruvate, 0.1% 2-mercaptoethanol and 1% P/S) supplemented with GM-CSF (PeproTech, #315-03), FLT-3L (PeproTech, #250-31L) and IL-6 (PeproTech, #216-16) according to the supplemental Table 1. After 10 days of expansion, BMDCs were collected by centrifugation, resuspended in the complete IMDM media and used for experiments.

For coculture experiments, CD4 or CD8 T cells were purified from the spleen of OT-II or OT-I mice respectively. Briefly, spleen was mechanically dissociated through a 100 μM cell strainer (Falcon^®^), cells were centrifuged and red blood cells were lysed. After washing, cells were isolated by negative magnetic sorting using a Dynabeads™ Untouched™ Mouse CD4 or CD8 Cells kit (Invitrogen™ #11415D #11417D) according to the manufacturer’s instructions. Cells were resuspended in complete RPMI media (10% FBS, 1% NEAA, 1% sodium pyruvate, 1% P/S) and used immediately.

### Lipid nanoparticle preparation and complexation with mRNA

Lipid nanoparticles (LNP) were prepared as described previously (36). Briefly, a lipid phase was prepared containing triglycerides (Suppocire NB, Gattefossé and super-refined soybean oil, Croda Uniqema) and phospholipids (Lipoid SPC3, Lipoid) supplemented with the cationic lipid DOTAP (1,2-dioleoyl-3-trimethylammonium-propane chloride, Avanti Polar Lipids) and fusogenic lipid DOPE (1,2-dioleoyl-sn-glycero-3-phosphoethanolamine, Avanti Polar Lipids). A second aqueous phase containing the PEGylated surfactant PEG-40 Stearate (Myrj S40, Croda Uniqema) was prepared in Phosphate-buffered saline (PBS) (#806552, Sigma). Both aqueous and lipid phases were mixed together through high-frequency sonication. Purification of LNP was performed by extensive dialysis in 100 volumes of LNP buffer: 154 mM NaCl, 10 mM HEPES, and pH 7.4 using endotoxin-free ultra-pure water (TMS-011-A, Sigma) and 12–14 kDa MW cut-off membranes (ZelluTrans/Roth T3). Lastly, the LNP solution was sterilized by filtrating through a 0.22-μm Millipore membrane under aseptic conditions and stored at 4°C until required.

LNP were complexed with Cherry, Egfp or Ova mRNA (Tebu bio #040L-7203-100; #040L-7601-100; #040L-7610-100) in IMDM media. The required volumes of mRNA and LNP were calculated for obtaining a N/P ratio of 6 (ratio of positively-chargeable polymer amine (N = nitrogen) groups to negatively-charged nucleic acid phosphate (P) groups). The LNP and mRNA were gently homogenized by pipetting and kept for 10 min at room temperature before immediate use for downstream experiments.

### Cell incubation with LNP-mRNA complexations and inhibitors

For BMDC treatment, 0.3 M cells were seeded in 96 well plates in 200 μL of IMDM media deprived in FBS and P/S and containing GM-CSF (5 ng/ml) and FLT-3L (25 ng/ml). After an overnight incubation at 37°C under 5% CO_2_, 20 μL of LNP, Ova mRNA or LNP-Ova mRNA complexation in IMDM was added in each corresponding well. 20 μL of IMDM media alone was used for the no treatment condition (vehicle). For some wells, after 6 hours of transfection the media was changed and the assessment of cytokines in the cell supernatant was performed at day 1.

For the experiments with TLR inhibitors, BMDCs were pretreated for 3 hours with either TLR4 inhibitor TAK-242 (InvivoGen, #tlr-cli95) at three different concentrations (1.25 μM, 2.5 μM, and 5 μM) or with the TLR8 inhibitor CU-CPT9a (2.5 μM, 5 μM, and 10 μM) (InvivoGen, #inh-cc9a) and incubated at 37 °C. For the experiments with the antioxidant drugs, the pretreatment with diphenyleneiodonium (DPI) (0.1 μM, 1 μM, and 10 μM), a ROS-dependent NOX inhibitor (Sigma-Aldrich, D2926) or with S3QEL2 (1 μM, 10 μM, and 30 μM), a mitochondrial ROS (mROS) inhibitor (Sigma-Aldrich, SML1554) lasted 1 hour. The cell supernatants were collected after 24 hours of LNP-Ova mRNA complex incubation.

For coculture experiments, 40 000 BMDCs treated with 125 ng of vectorized mRNA for 6 hours were washed with IMDM media and 200 μL of CD4 or CD8 T cells at 550 000 cells/mL were added. Cocultures were incubated for 48 hours at 37°C under 5% CO_2_ before supernatant collection and cell analysis. Protein Transport Inhibitor cocktail (PTI, eBioscience) was added for the last 6 hours of culture.

### Murine tumor models and *in vivo* treatments

B16OVA (2 × 10^5^) or E.G7-OVA (4 × 10^5^) cells were intradermally (i.d.) implanted into the flank of female C57BL/6 mice in a final volume of 100 μL of sterile phosphate buffered saline (PBS). Digital calipers were used to measure the perpendicular diameters of the tumors three times a week. In agreement with ethical guidelines, mice were euthanized when the tumor size reached 250 mm^2^ or when tumor ulcerations were visible.

#### Preventive therapy

3 μg of vectorized Cherry or Ova mRNA prepared in IMDM were injected intraperitoneally (i.p.) (100 μL per mouse) at day 0, 3 and 8. At day 14, B16OVA cells were inoculated and mice were sacrificed at day 30 for subsequent analysis. For long term experiment, B16OVA were grafted at day 49.

#### Curative therapy

After B16OVA or E.G7-OVA tumor engraftment (day 0), mice were injected i.p. with 3 μg of vectorized Cherry, Egfp or Ova mRNA prepared in IMDM at the intervals indicated in the figure legends and sacrificed at day 17.

For CD8 or NK cell depletions, anti-CD8β (clone 53-5.8, BioXCell) or anti-NK1.1 (clone PK136, BioXCell) or their respective isotype control antibody (clone HRPN or C1.18.4, BioXCell) was injected i.p. at 150 μg per mouse at day 2, 7 and 13 after B16OVA cells engraftment. For PD-1 blockade, anti-PD-1 (clone RMP1-14, BioXCell) or its isotype control antibody (clone 2A3, BioXCell) was injected i.p. at 100 μg per mouse at day 7, 12 and 17 relative to B16OVA cell inoculation. Immunizations with 3 μg of vectorized mRNA were performed at day 3, 6 and 9.

#### Cancer challenge

To challenge survival mice after tumor regression, a new B16OVA or B16F10 tumor was engrafted in the contralateral site (2 × 10^5^ cells in a final volume of 100 μL of sterile PBS).

### Cytokine measurement

Cell culture supernatant or serum were analyzed by ELISA for IL-6 (#88-7064, Invitrogen), TNFα (#88-7324, Invitrogen) or IFNγ (#88-8314, Invitrogen) according to the manufacturer’s instructions.

### Toxicity assessment

Toxicity was evaluated by measuring the lactate dehydrogenase (LDH) level in cell supernatants using the CytoTox-ONE™ Homogeneous Membrane Integrity Assay kit (Promega, G7892) according to the manufacturer’s instructions. Briefly, cell lysis was induced with a lysis buffer and was used as positive control for LDH release. 50 μl of cell supernatant were dropped in a flat bottom 96-well plate, incubated with 50 μl of CytoTox-ONE™ reagent for 30 seconds on the microplate shaker and for 20 minutes in the dark at room temperature (RT). A stop solution (25 μl) was added to each well and the plate was placed on the microplate shaker for 10 seconds. Ultimately, the fluorescence was measured at an excitation wavelength of 560 nm and an emission wavelength of 590 nm using a CLARIOstar^®^ microplate reader (BMG LABTECH).

### Flow cytometry

Tumors were mechanically dissociated with scissors and digested enzymatically using the Tumor Dissociation Kit mouse and the gentleMACS Dissociator (Miltenyi Biotec) according to the manufacturer’s instructions. After filtration through a 70 μm cell strainer (Miltenyi Biotec), the single-cell suspension was washed two times with complete RPMI-1640. Following the last centrifugation, the pellet was resuspended with a volume of complete RPMI-1640 to obtain a final concentration of 14 mg of tumor for 100 μL. Then, 100 μL of cell suspension was dropped in U 96-well plates for assessment of immune cell populations in the same amount of tumor for each one. Spleens and tumor draining lymph nodes (TDLN) were mechanically dissociated through 100 μm cell strainers (Falcon^®^). After red blood cell lysis, splenocytes were washed one time with complete RPMI-1640. 1 million of splenocytes and 0.5 million of cells from TDLN were used for staining in U 96-well plates. After PBS 1X washing, cells were stained with LIVE/DEAD™ Fixable Red Dead Cell Stain Kit for 30 min protected from light at RT. Fc receptors were blocked with anti-CD16/CD32 (TruStain FcX™; Biolegend) and cells were stained for cell surface antigens. For intracellular staining, cells were fixed and permeabilized using the Foxp3 / Transcription Factor Staining Buffer Set (eBioscience™). For *ex vivo* stimulation, intratumoral immune cells and splenocytes were restimulated with class II-restricted OVA-peptide (323-339) and/or class I-restricted OVA-peptide (257-264) for 4 hours or 16 hours respectively, both with PTI. Events were acquired on BD-LSRII flow cytometer (BD Biosciences, Le Pont-De-Claix, France), collected with BD FACSDiva 6.3.1 software and analyzed using Flowlogic software (Miltenyi Biotec). All antibodies used for extracellular and intracellular stainings are listed in the supplemental table 2.

### Immunohistochemistry

FFPE blocks were sectioned at 4 μm thickness using a microtome. Paraffin was removed and antigen retrieval was performed using a PT-Link apparatus (Agilent) at 97°C for 20 minutes. After these steps, endogenous peroxidases were blocked for 5 minutes (Agilent, ref. SM801) and then the slides were incubated with CD8 antibody (Histosure, polyclonal, 1/100, 30 minutes), CD4 antibody (Histosure, clone Rb78H9D2, 1/900, 30 minutes) or PD1 antibody (Cell Signaling, clone D7D5W, 1/50, 60 minutes). Slides were washed and then incubated with secondary HRP polymer reagents (Vector Laboratories, ref. MP7451) for 20 minutes. The slides were then washed and incubated with magenta chromogen (Agilent, ref. GV92511-2) for 5 minutes. After a final washing step, the slides were finally counterstained with haematoxylin (Agilent, ref. SM806) for 5 minutes. All staining steps were performed using an Autostainer AS48 (Agilent). Slides were digitized using a VS200 slide scanner (Evident) at X20 magnification. Analysis of staining and quantification was performed using QuPath software (Bankhead, 2017, Sci reports).

### Bulk RNA-Sequencing from B16OVA tumors

Tumors were snap frozen in liquid nitrogen and stored at -80°C until use. RNA extraction, library preparation, RNA sequencing (RNA-seq), and determination of differentially expressed genes were performed by BGI genomics (Hong Kong, China). Total RNA was extracted from frozen B16OVA tumors lysed by TissueLyser II (QIAGEN) using phenol-chloroform extraction method with Trizol (Invitrogen) and Isopropanol (XILONG). The quality of the total RNA and the sample RNA concentration were determined by using the Bioanalyzer 2100 (Agilent). 1 μg of total RNA was used to build mRNA libraries using MGIEasy RNA Library Prep kit (MGI, China) following the manufacturers’ instructions. Briefly, mRNAs were first enriched using oligo(dT)-attached magnetic beads. Following mRNA fragmentation, first and second strand cDNA was synthesized using random primers (MGI, China). cDNA fragments were end-repaired, adenylated at 3⍰ ends, purified with magnetic beads by using a linker which was connected to the A base under enzymatic reaction, and enriched by PCR. After controlling the quality of PCR products on the Bioanalyzer 2100 (Agilent Technologies Inc.), double stranded PCR products were heat denatured and circularized by the splint oligo sequence. The single strand circle DNA (ssCir DNA) were formatted as the final library. The library was amplified to make DNA nanoball (DNB) which had more than 300 copies. The DNBs were load into the patterned nanoarray and sequenced on the DNBseq G400 (MGI, China) with read length as 2×150 bp paired end reads. Clean reads were then aligned against Mus musculus mm39 reference genome (source: UCSC) using HISAT. Analysis were performed with Dr. Tom online system and details are mentioned in the figure legends.

### Statistical analysis

Data were analyzed with Prism software version 8.0.2 (Graph Pad software, La Jolla, California, USA) and shown as mean ± SEM. Data normality was tested with a D’Agostino and Pearson test. Mann–Whitney U test were used for comparisons of two groups. For multiple comparisons one-way or two-way ANOVA with Bonferroni’s or Dunnett’s multiple comparison post-hoc tests or Kruskal-Wallis test with Dunn’s multiple comparisons test were performed when appropriate. For survival curve statistics, log-rank (Mantel-Cox) test was used. For correlation analysis, a Pearson r correlation coefficient was calculated. Statistical test used are specified in each figure legends. *P* < 0.05 was considered statistically significant.

## Results

### Activation of DCs treated with LNP-Ova mRNA and ability of treated-DCs to specifically activate T cells *in vitro*

Given the crucial role of DCs as antigen presenting cells (APCs) in the antitumor effect of the mRNA-based vaccine, we tested whether our vaccinal strategy has an impact on DC activation *in vitro*. For this, we tested if Ova mRNA alone or vectorized affects the secretion of proinflammatory cytokines in BMDCs for a short and a long transfection timing. While LNP or Ova mRNA alone does not lead to IL-6 or TNFα secretion, Ova mRNA vectorized with LNP for 6h and 24h of transfection increases the concentrations of these two cytokines into the DC supernatant at day 1 (**figure 1A,B**). Previous studies reported that mRNA-based vaccines activate TLR pathways (24,25) and TLR activation increases ROS production in DCs (44). We thus evaluated the contribution of TLR4, TLR8 and ROS signaling in mediating the proinflammatory effect of the vectorized Ova mRNA in BMDCs by using specific inhibitors. The inhibition of TLR4 and TLR8 by TAK-242 and CU-CPT9a, respectively, drives a drastic decrease in the LNP-Ova mRNA-mediated IL-6 release (**figure 1C,D**). Likewise, the inhibition of NOX-dependent ROS and mitochondrial ROS (mROS) by DPI and S3QEL 2, respectively, reduces the IL-6 secretion in response to vectorized mRNA (**figure 1E,F**). Importantly, for all the doses, the TLR and ROS inhibitors used in combination with the LNP-Ova mRNA complex do not exhibit any effect on cell viability excluding cell death as a potential event leading to the lower IL-6 secretion (**supplemental figure 1A,B**).

**Figure 1:**
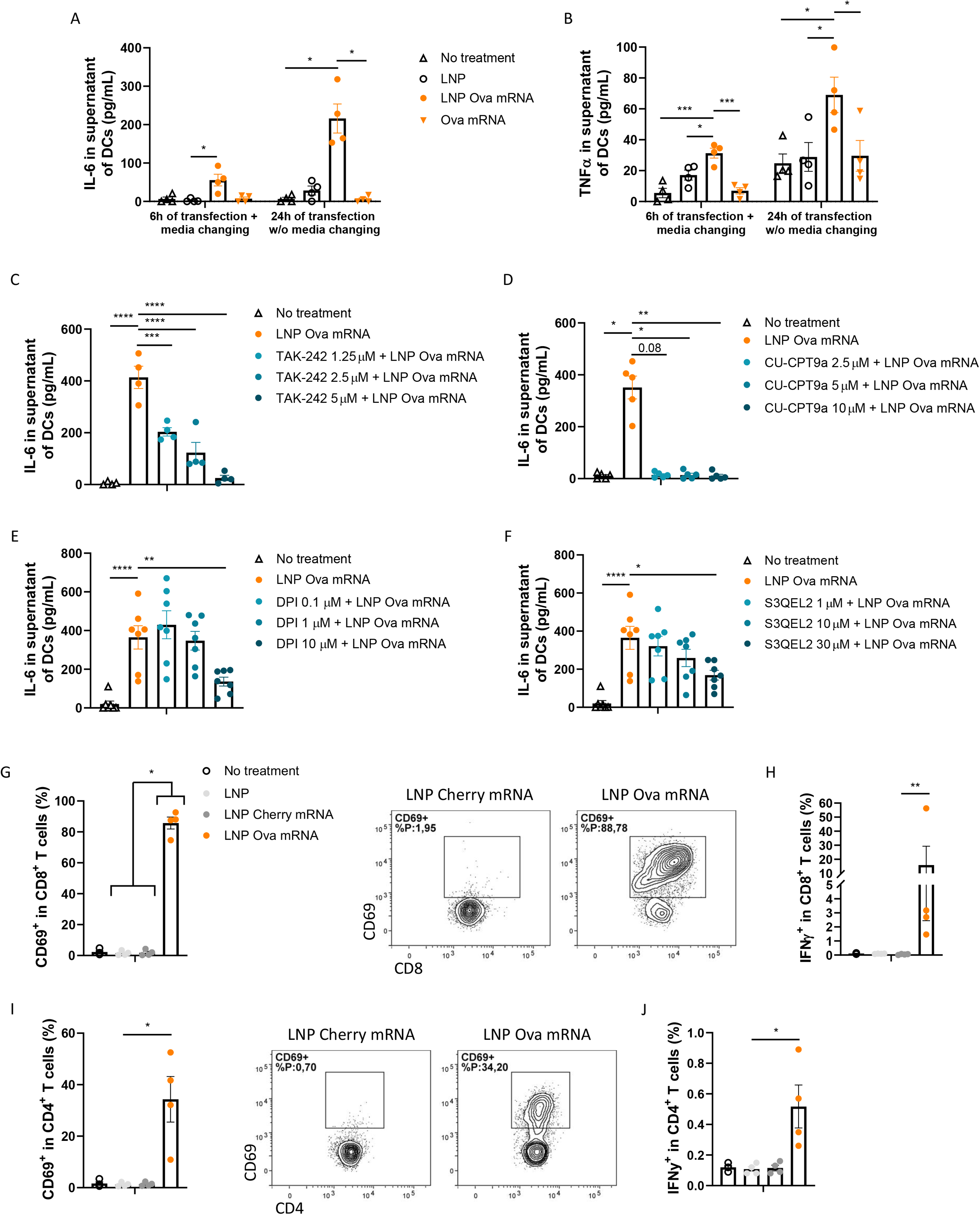
LNP-Ova mRNA mediates BMDC and T cell activation *in vitro*. **(A**,**B)** IL-6 (A) and TNFα (B) concentration in BMDC supernatant at day 1 following a 6h or 24h time incubation with vehicle, LNP, LNP-Ova mRNA or Ova mRNA. **(C-F)** IL-6 concentration in BMDC supernatant following 3 hours of pretreatment with TLR4 inhibitor TAK-242 at three different concentrations (1.25 μM, 2.5 μM, and 5 μM) (C) or TLR8 inhibitor CU-CPT9a (2.5 μM, 5 μM, and 10 μM) (D) or following 1 hour of pretreatment with ROS-dependent NOX inhibitor DPI (0.1 μM, 1 μM, and 10 μM) (E) or mitochondrial ROS (mROS) inhibitor S3QEL2 (1 μM, 10 μM, and 30 μM) (F) and 24h of LNP-Ova mRNA incubation. Mean ± SEM of 4 to 7 independent experiments where each dot represents the mean of 2 replicates. **(G-J)** CD8^+^ T cells (G,H) or CD4^+^ T cells (I,J) from OT-I or OT-II spleen respectively were cocultured for 48 hours with BMDCs treated with vehicle, LNP, LNP-cherry mRNA or LNP-Ova mRNA. Frequencies of CD69^+^ (G,I) and IFNγ^+^ (H,J) were assessed by flow cytometry. Mean ± SEM of four independent experiments where each dot represents the mean of 2 to 3 replicates. *P* values (\**p* < 0.05, \*\**p* < 0.01, \*\*\**p* < 0.001, \*\*\*\**p* < 0.0001) determined by Kruskal-Wallis test with Dunn’s post hoc testing (A,D,G-J) or one-way ANOVA with Dunnett’s post hoc testing (B,C,E,F).

To evaluate if BMDCs treated with LNP-Ova mRNA complex are able to translate mRNA, process antigens and present them to T cells, we cocultured them with CD8^+^ or CD4^+^ T cells isolated from spleen of OT-I or OT-II mice, respectively. The frequency of activated CD8^+^ lymphocytes (CD69^+^) is considerably increased (**figure 1G**), together with an enhanced production and secretion of IFNγ (**figure 1H; supplemental figure 1C**), TNFα (**supplemental figure 1D,E**) and IL-2 (**supplemental figure 1F**) compared with control conditions. Although to a lesser extent, similar results for CD69, IFNγ and TNFα are observed in CD4^+^ T cells (**figure 1I,J; supplemental figure 1G-I**).

Altogether, these results indicate that LNP-Ova mRNA activates DCs *in vitro* in a TLR4/8 and ROS dependent manner, effectively delivers antigen-encoding mRNA to BMDCs and promotes efficient mRNA translation and antigen presentation for specific CD8^+^ and CD4^+^ T cell activation.

### Antitumor efficacy of LNP-Ova mRNA-based vaccine in a cancer prevention setting

Because of the capacity of the LNP-Ova mRNA complex to lead to a cellular immune response *in vitro* we next evaluate whether vectorized Ova mRNA prevents tumor growth *in vivo*. For this purpose, we immunized healthy adult mice three times with 3 μg of vectorized mRNA, grafted B16OVA melanoma cells intradermally 6 days after the last immunization and followed the tumor size evolution (**figure 2A**). In the LNP-Ova mRNA vaccine group, the tumor growth was completely impeded (6/6 mice) in comparison with the control group vaccinated with a mRNA encoding an irrelevant protein (**figure 2B,C**). Then, we assessed the activity of T lymphocytes at the periphery level since no tumors were available in the treated group for subsequent analysis. Under LNP-Ova mRNA treatment conditions, the frequency of CD8^+^ T cells from the spleen producing IL-2, IFNγ, GzmB or TNFα, coproducing IFNγ and TNFα and expressing PD-1 increases when compared to the control group (**figure 2D-I**) based on fluorescence minus one (FMO) controls for positive threshold determination (**supplemental figure 2A**). Similar changes in CD4^+^ T cells from the spleen, CD8^+^ and CD4^+^ T cells from the TDLN producing IL-2, IFNγ and TNFα were not detected (**supplemental figure 2B-J**). Moreover, we observed a higher concentration level of IFNγ in the splenocyte supernatant of the LNP-Ova mRNA group which would reflect the activation of the CD8^+^ rather than the CD4^+^ T cell, consistent with the flow cytometry results (**supplemental figure 2K**). Finally, we assessed whether the vaccine-induced antitumoral effect is still efficient even after a longer time (41 days) between the last vaccination and the tumor cell inoculation. Immunization with LNP-Ova mRNA leads to complete protection upon tumor challenge also with this experimental setting (**supplemental figure 2L,M**). Overall, these results demonstrate that vectorized Ova mRNA can be used as an effective vaccine to prevent tumor development from OVA-expressing cancer cell engraftment together with the establishment of activatable and potentially mobilizable CD8^+^ T cells from the periphery.

**Figure 2:**
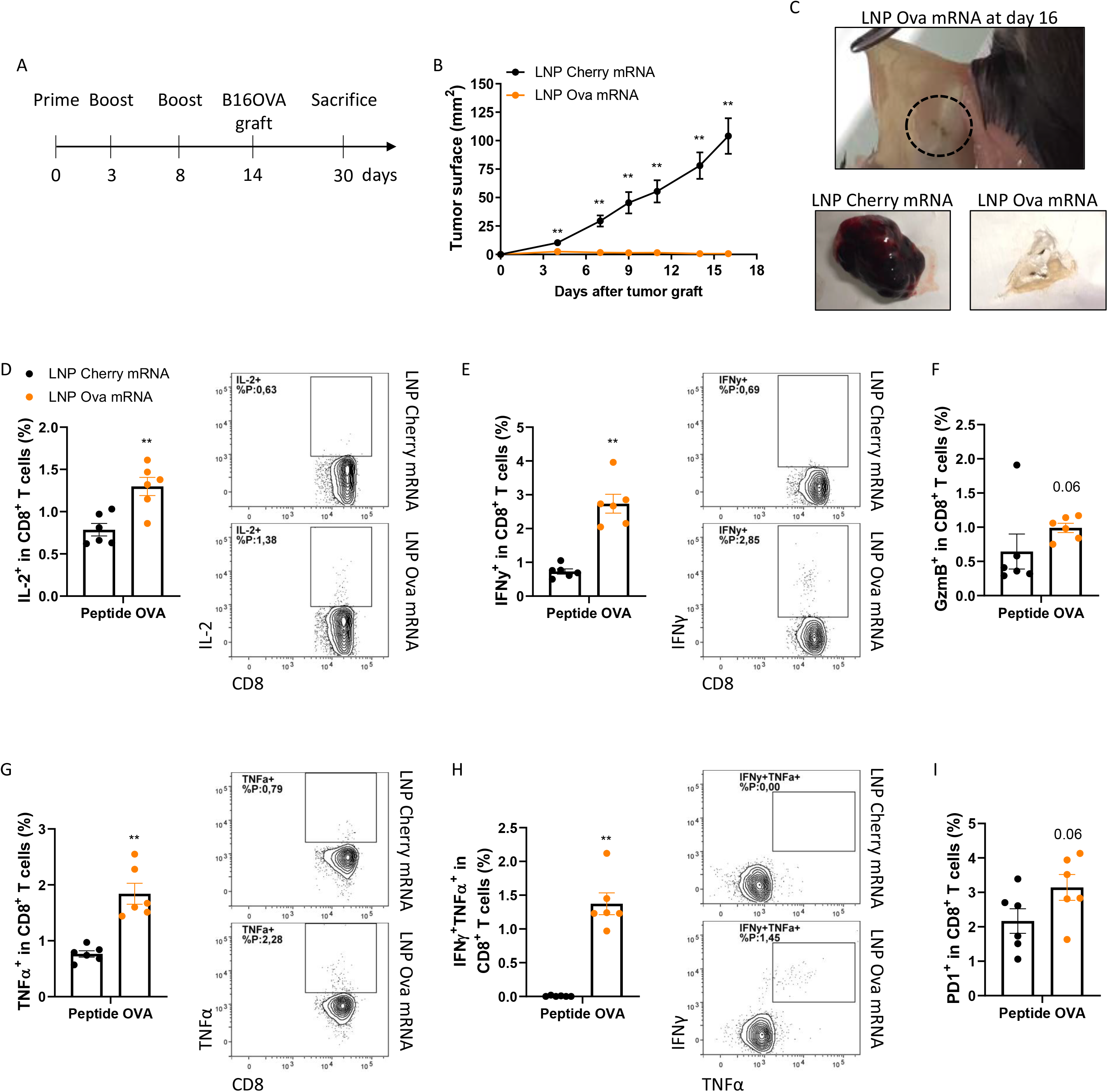
Antitumor effect of LNP-Ova mRNA complex as a preventive vaccine. **(A-C)** Tumor growth in LNP-Ova mRNA immunized mice injected i.d. with B16OVA melanoma cells. LNP-Cherry mRNA was used for the control group (B). Representative pictures of the B16OVA tumor from LNP-Ova mRNA or LNP-Cherry mRNA treated mice at sacrifice (C). **(D-I)** CD8^+^ T cell analysis from spleen of B16OVA tumor-bearing mice immunized with LNP-Ova mRNA or LNP-Cherry mRNA. Following *ex vivo* restimulation with OVA peptides, IL-2 (D), IFNγ (E), GzmB (F), TNFα (G) production, IFNγ and TNFα coproduction (H) and PD-1 expression (I) were assessed by flow cytometry. Mean ± SEM of *n* = 6 mice per group. *P* values (\**p* < 0.05, \*\**p* < 0.01, \*\*\**p* < 0.001, \*\*\*\**p* < 0.0001) determined by Mann-Whitney test (B-I).

### Antitumor efficacy of LNP-Ova mRNA-based vaccine in a cancer therapeutic setting

We tested the therapeutic efficacy of our vaccinal strategy using two OVA-expressing murine tumor models. After B16OVA or E.G7-OVA tumor cell inoculation, three immunizations with 3 μg of vectorized Ova mRNA were performed (**supplemental figure 3A**,**B**). In the groups treated with LNP-Ova mRNA, regardless of the type of tumor tested, tumor growth and weight at sacrifice are drastically reduced while spleen weight is not affected when compared to the control groups (**figure 3A,B; supplemental figure 3C,D**). Intratumoral T-cell specific immune response was assessed by flow cytometry after restimulation with OVA peptides for 4h00 (for gating strategy see **supplemental figure 3E**) and immunohistochemistry (IHC). LNP-Ova mRNA-based vaccination increases the percentage of CD45^+^ cells in live cells, CD8^+^ T cells in CD45^+^ cells as well as the number of B16OVA tumorinfiltrating CD45^+^, CD8^+^ and CD4^+^ cells compared with the LNP-control mRNA-treated mice (**figure 3C-F; supplemental figure 3F-H**). In addition, we noted higher frequencies of intratumoral activated CD8^+^ T cells expressing PD-1, CD69, producing IFNγ, TNFα or GzmB as well as coproducing IFNγ and TNFα or IFNγ and GzmB in the LNP-Ova mRNA group (**figure 3G-J; supplemental figure 3I-L**). Concerning the CD4^+^ T cells, only the frequencies of CD69^+^ and TNFα^+^ were significantly increased in this group (**supplemental figure 3M-P**). For the E.G7-OVA tumor model, similar results were observed. The LPN-Ova mRNA-based vaccination leads to a higher frequency of intratumoral CD8^+^ T cells in CD45^+^ cells as well as higher frequencies of CD8^+^ T cells expressing PD-1 and coproducing cytokines and GzmB when compared to the control group (**supplemental figure 3Q-T**). Although the frequency of CD4^+^ T cells in CD45^+^ cells was not affected, the frequency of CD4^+^ T cells producing IFNγ and TNFα were increased in the LNP-Ova mRNA group (**supplemental figure 3U-X**). We also observe a negative correlation between percentage of activated CD8^+^CD69^+^ T cells and tumor weight at sacrifice which is more significant than with activated CD4^+^CD69^+^T cells (**figure 3K**). Because of the massive CD8^+^ T cell infiltration into the tumor, their clear enhanced functional profile and their correlation with the tumor weight, we completed our study by depleting CD8^+^ T in the B16OVA tumor model. This experiment reveals that LNP-Ova mRNA-based vaccination efficacy is CD8^+^ T cell-dependent since the median survival of 34.5 days in this group was reduced to 21.5 days in the group vaccinated with LNP-Ova mRNA and blocked for CD8^+^ T cells, a value close to the control group (20.5 days) (**figure 3L; supplemental figure 3Y,Z**). Overall, these results indicate that vectorized Ova mRNA is an effective vaccine strategy for inhibiting tumor growth in a curative setting which mainly relies on enhanced CD8^+^ T cell effector function.

**Figure 3:**
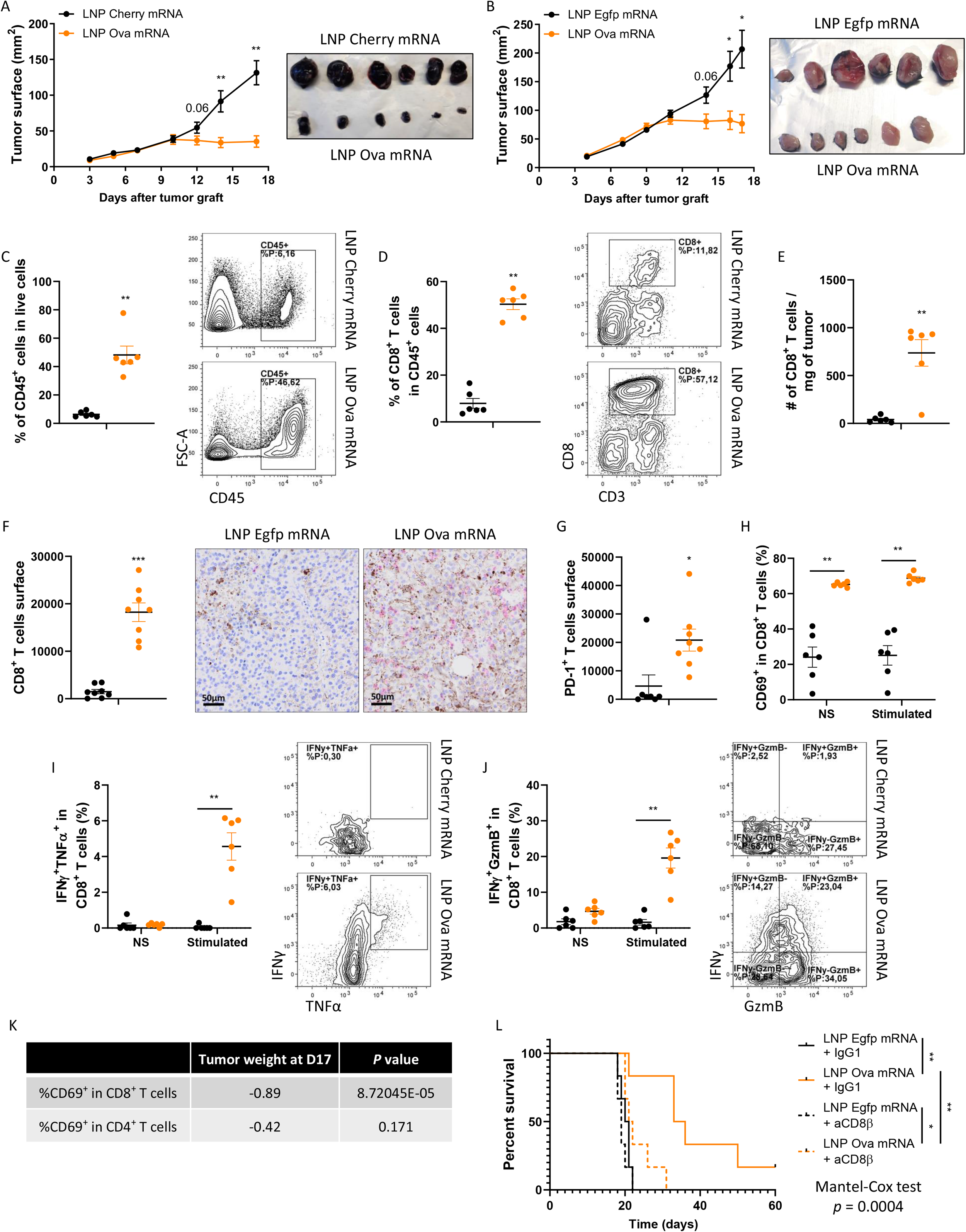
Antitumor effect of LNP-Ova mRNA complex as a therapeutic vaccine. **(A**,**B)** Tumor growth in mice injected with B16OVA melanoma cells (A) or E.G7-OVA lymphoma cells (B) and then treated with LNP-Ova mRNA or LNP-irrelevant mRNA. **(C-J)** Intratumoral frequency of CD45^+^ cells in live cells (C), CD8^+^ T cells in CD45^+^ cells (D), number of CD8^+^ T cells per mg of B16OVA tumor tissue (E) from LNP-Ova mRNA or LNP-Cherry mRNA-treated mice. Surface of CD8^+^ T cells (F) and PD-1^+^ T cells (G) analyzed by IHC for each tumor group. Intratumoral frequency of CD8^+^ T cells expressing CD69 (H) and coexpressing IFNγ and TNFα (I) or IFNγ and GzmB (J) from B16OVA tumor-bearing mice treated with LNP-Ova mRNA or LNP-irrelevant mRNA following OVA peptide stimulation. **(K)** Correlations between intratumoral CD69^+^CD8^+^ T cells frequency or CD69^+^CD4^+^ T cells frequency and tumor weight at day 17 in B16OVA-bearing tumor mice treated with LNP-Ova mRNA or LNP-Cherry mRNA. **(L)** Survival in B16OVA tumor-bearing mice treated i.p. with LNP-Egfp mRNA or LNP-Ova mRNA as well as with anti-CD8β or its control IgG1. Mean ± SEM of n = 6 mice per group. P values (\**p* < 0.05, \*\**p* < 0.01, \*\*\**p* < 0.001, \*\*\*\**p* < 0.0001) determined by Mann-Whitney test (A-J) or Pearson r correlation test (K) or log-rank (Mantel-Cox) test (L).

### LNP-Ova mRNA-based vaccine reprograms the intratumoral immune-related transcriptional signature in a favorable way to anti-PD-1 response

An unbiased RNA sequencing (RNA-seq) was performed following two immunizations at day 9 to evaluate early changes in gene expression induced by LNP-Ova mRNA-based vaccine in B16OVA tumor bearing mice (**supplemental figure 4A**). The results reveal the upregulation of many genes involved in the immune response as shown in the volcano plot, especially genes related with tumoricidal effector T cells (*Ifng, Gzma, Gzmb, Cd8a, Cd69*) and lymphocyte attraction (*Cxcl10, Cxcl11, Cxcl9*) (**figure 4A**). Overall, 487 genes were differentially expressed after vaccine treatment (defined as a log2 fold change > 2 and *P* < 0.005) (**supplemental figure 4B**). Gene Set Enrichment Analysis (GSEA) based on these 487 genes shows that LNP-Ova mRNA treatment leads to a selective enrichment of biological pathways involving notably the “cellular response to IFN-beta” and “cellular response to IFN-gamma” (**figure 4B-D**). Then, a KEGG pathway classification was realized and the higher number of genes modified (123) in response to the LNP-Ova mRNA-based vaccine referred to the “immune system” pathway (**figure 4E**). Based on these 123 genes, we performed a KEGG pathway relationship network (**supplemental figure 4C**) and selected, ranked by the number of genes and *p* value, the three most significative pathways to generate gene expression heatmaps which are related to the “T cell receptor signaling pathway”, “chemokine signaling pathway” and “antigen processing and presentation” (**figure 4F, G** and **supplemental figure 4D**, respectively). Importantly, our vaccinal strategy converted the cold intratumoral gene signature to an immune-related signature correlating with clinical benefit to PD-1 blockade in melanoma patients (8) and potentially predicting the synergetic action of this therapeutic combination (**supplemental figure 4E**). Each gene of all the explored pathways here shows a significant upregulation in the LNP-Ova mRNA treated group (log2 fold change > 2 and *P* < 0.001). Altogether, these proinflammatory gene expressions cooperate to build up an immunoreactive tumor microenvironment that favored CD8^+^ T cell accumulation and activation in a favorable way to anti-PD-1 response.

**Figure 4:**
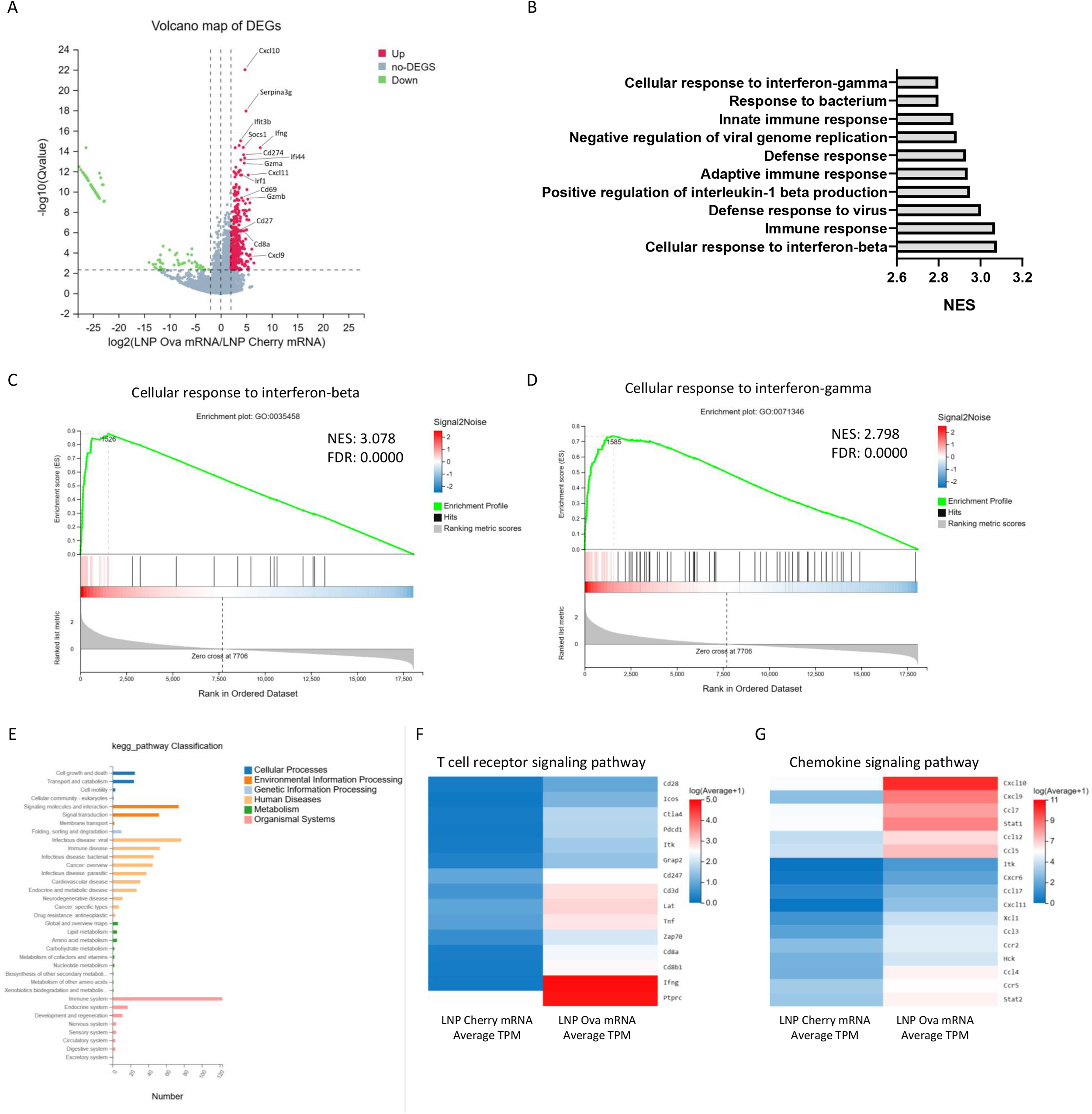
LNP-Ova mRNA vaccine produces an inflamed immune gene signature. Mice bearing B16OVA tumors were treated with LNP-Cherry mRNA or LNP-Ova mRNA twice and analyzed by RNA-seq. **(A)** Vulcano plot showing the fold change (log2, x-axis) and statistical significance (−log10 Q value, y-axis) of differentially expressed genes (DEGs) (log2 fold change > 2 and *P* < 0.005). **(B)** Top 10 of selective enrichment of biological pathways following GSEA based on 487 DEGs between LNP-Cherry mRNA and LNP-Ova mRNA groups (log2 fold change > 2 and *P* < 0.005). **(C**,**D)** Enrichment plot of indicated signatures. Normalized Enrichment score (NES), False Discovery Rate (FDR) q value shown for the two gene sets. **(E)** KEGG pathway classification based on 487 DEGs between LNP-Cherry mRNA and LNP-Ova mRNA groups (log2 fold change > 2 and *P* < 0.005). **(F**,**G)** Heatmaps of genes related to “T cell receptor signaling pathway” **(F)** and “chemokine signaling pathway” **(G)** following the selection of the most significative pathways from a KEGG pathway relationship network based on the 123 genes related to the “immune system” pathway. N = 4 mice per group.

### Anti-PD-1 blocking antibody synergizes with anticancer effect of LNP-Ova mRNA-based vaccine

Given that our vaccine strategy renders the immunologically inactive and anti-PD-1-resistant B16OVA tumor model immunologically active, we tested whether LNP-Ova mRNA treatment would result in sensitization of this model to anti-PD-1 immunotherapy. Furthermore, in addition to an increased frequency of CD8^+^ T cells expressing PD-1 in B16OVA and E.G7-OVA tumor models in response to the LNP-Ova mRNA treatment (**figure 3G; supplemental figure 3R**), we observed a greater fraction of inflammatory monocytes that expresses PD-L1 in the E.G7-OVA lymphoma model (**supplemental figure 5A-C**) as well as a higher gene expression of *Cd274* and *Pdcd1* in B16OVA tumor (**figure 4A, F**). These results suggest that the PD-1/PD-L1 axis can be a possible mechanism of immune escape in response to long-term vaccine-induced antitumor action. To test this, we immunized mice with LNP-Ova mRNA and treated them with anti-PD-1 using the B16OVA tumor model (for treatment schedule see **supplemental figure 5D**). In these experiments we still show the vaccine-mediated antitumor effect, and, we observe that this effect is transient. Indeed, the resumption of tumor growth appears in majority of mice; only 3/12 mice survive after 100 days following tumor cell injection in the LNP-Ova mRNA-treated group (**figure 5A,B; supplemental figure 5E**), whereas the combination with anti-PD-1 considerably improves the tumor growth control and the mouse survival since 9/12 mice are still alive at day 100 in the group treated with Ova mRNA-based vaccine and anti-PD-1 (**figure 5A,B; supplemental figure 5E**). In order to better understand the potential early immune-related mechanisms underlying the beneficial effect of this combinatorial treatment, we analyzed the intratumoral myeloid, T and NK cells at day 9, two days after the first injection of anti-PD-1. At this timing we mainly observed a vaccine effect on myeloid cells leading to a higher intratumoral frequency of activated DCs, activated macrophages, inflammatory monocytes (*p* < 0.0001, *p* = 0.02, *p* < 0.0001 respectively) and a reduced fraction of M2 CD206^+^ macrophages (p < 0.0001) (**figure 5C-F**; for gating strategy see **supplemental figure 5F**). While a trend was notable for macrophages, DCs and inflammatory monocytes show a statistically significant more intensive expression of PD-L1 in LNP-Ova mRNA treated mice especially in the group combined with anti-PD-1 (**figure 5G-I**). Interestingly, while the vaccine treatment raises the fraction of CD8^+^ T cells expressing GzmB, IFNγ and PD-1 (**figure 5J,K**; for gating strategy see **supplemental figure 5G**), we observe a close statistically significative interaction between aPD-1 and vaccine therapies leading to a partial prevention of PD-1^+^TIM-3^+^ CD8^+^ T cell accumulation and a statistically significative interaction leading to a higher proportion of CD4^+^ T producing GzmB in the group treated with the treatment combination (**figure 5L,M**). The IFNγ serum concentration is also higher in the groups treated with the vaccine in comparison to the LNP-Egpf mRNA + IgG2a group more probably related to the CD8^+^ T cell-mediated IFNγ secretion since the production of CD4^+^ T cells was not modulated by any treatment at this time point (**supplemental figure 5H,I**). Last, although the LNP-Ova mRNA treatment enhances the intratumoral frequency of NK cells and their production of GzmB, the blocking of these cells does not impair the vaccine-mediated antitumoral efficacy while it shortens the survival of mice treated with LNP-Egpf mRNA (**supplemental figure 5J-L**). Altogether, these results highlight the apparition of a PD-1 dependent immune escape process in response to the vaccine that can be block for improving the mice survival. At this early timing the vaccine and anti-PD-1 combination targets CD4^+^ T cells more particularly by enhancing their cytotoxic function.

**Figure 5:**
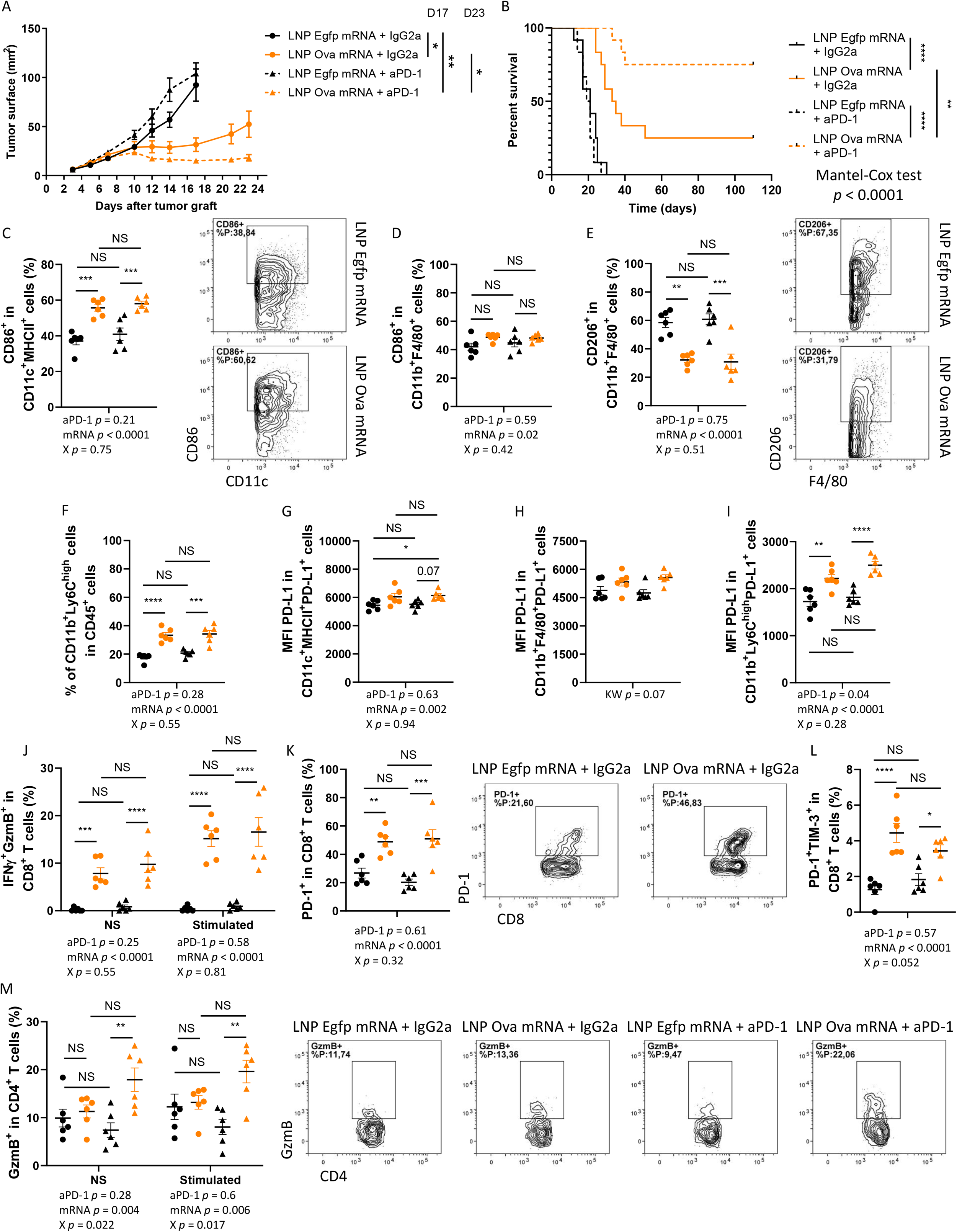
LNP-Ova mRNA vaccine cooperates with anti-PD-1 to suppress B16OVA tumor growth. **(A**,**B)** Tumor growth (A) and survival (B) in mice injected with B16OVA tumor cells and treated i.p. with LNP-Ova mRNA or LNP-Egfp mRNA as well as with anti-PD-1 or its control IgG2a. Mean ± SEM of n = 12 mice per group. P values (*p < 0.05, **p < 0.01, ****p < 0.0001) determined by Kruskal-Wallis test with Dunn’s post hoc testing (A, D17) and Mann-Whitney test (A, D23) or log-rank (Mantel-Cox) test (B). **(C-M)** Intratumoral frequency of CD86^+^ DCs (CD11c^+^MHCII^+^) (C), CD86^+^ macrophages (CD11b^+^F4/80^+^) (D), CD206^+^ macrophages (E) and proinflammatory monocytes (CD11b^+^Ly6C^high^) (F) PD-L1 expression (MFI) in PD-L1^+^ DCs (G), PD-L1^+^ macrophages (H) and PD-L1^+^ inflammatory monocytes (CD11b^+^Ly6C^high^) (I); intratumoral frequency of CD8^+^ T cells coexpressing IFNγ and GzmB (J), expressing PD-1 (K), coexpressing PD-1 and TIM-3 (L) and intratumoral frequency of CD4^+^ T cells expressing GzmB (M) from B16OVA tumor-bearing mice treated with LNP-Ova mRNA or LNP-Egfp mRNA as well as with anti-PD-1 or IgG2a. Mean ± SEM of n = 6 mice per group. P values (\**p* < 0.05, \*\**p* < 0.01, \*\*\**p* < 0.001, \*\*\*\**p* < 0.0001) determined by two-way ANOVA with Tukey’s post hoc testing (C-F,H-L) or Kruskal-Wallis test with Dunn’s post hoc testing (G).

### Long-term memory immune response in cancer survivor mice

In order to evaluate the memory antitumoral immune response and its ability to prevent the development of a new tumor, the surviving mice treated with vectorized Ova mRNA or the combination vectorized Ova mRNA and anti-PD-1 were challenged with a second injection of B16OVA at 103 days. Tumor growth in the challenged mice is largely blocked; the size of the tumors at day 5 of follow-up is already significantly lower than the values of the control group (**figure 6A**). Since the tumors were too small for analyzing the intratumoral immune response, we evaluated different immune cell populations in the spleen. Following an *ex vivo* restimulation, the production of IFNγ, TNFα or both (frequencies and MFI) by CD8^+^ splenocytes is higher in surviving mice than in animals of the control group (**figure 6B-F**). Similarly, Ova-specific CD8^+^ T cells (Ova dextramer) are more frequent in the spleens of challenged mice (**figure 6G**). Moreover, although the frequencies of DCs and B cells are not modified in CD45^+^ cells (**supplemental figure 6A-C**), their level of activation is higher (↑ MFI of CD86 in CD86^+^ DCs and ↑ frequency of CD86^+^MHCII^+^ B cells) in the spleen of treated mice having survived their 1^st^ tumor without PD-L1 expression changes (**figure 6H,I; supplemental figure E-G**). Finally, we challenged another batch of surviving mice with B16F10 cells in order to assess if they developed an antitumor memory immune response against antigens other than those derived from OVA. Although the tumor growth in the challenge group was not completely stopped as we observed with OVA-expressing B16 melanoma cells, the tumor size from day 4 until day 14 as well as the tumor weight at sacrifice were lower than in the control group (**figure 6J,K**). While the frequency of intratumoral activated CD8^+^ T cells was not different between these two groups (**supplemental figure 6H,I**), the frequency of activated NK cells was increased in the challenge group and negatively correlated with the final tumor weight (**figure 6L,M**). Altogether these results demonstrate that in responsive mice to our vaccinal strategy in combination or not with anti-PD-1 a long-lasting memory antitumoral immune response was induced in association with the peripheral establishment of polyfunctional and antigen-specific CD8^+^ T cells and activated APCs without PD-L1 expression-related immunosuppression. This memory immune response is not restricted to OVA-derived antigens suggesting that treatment-related antigen spreading occurs.

**Figure 6:**
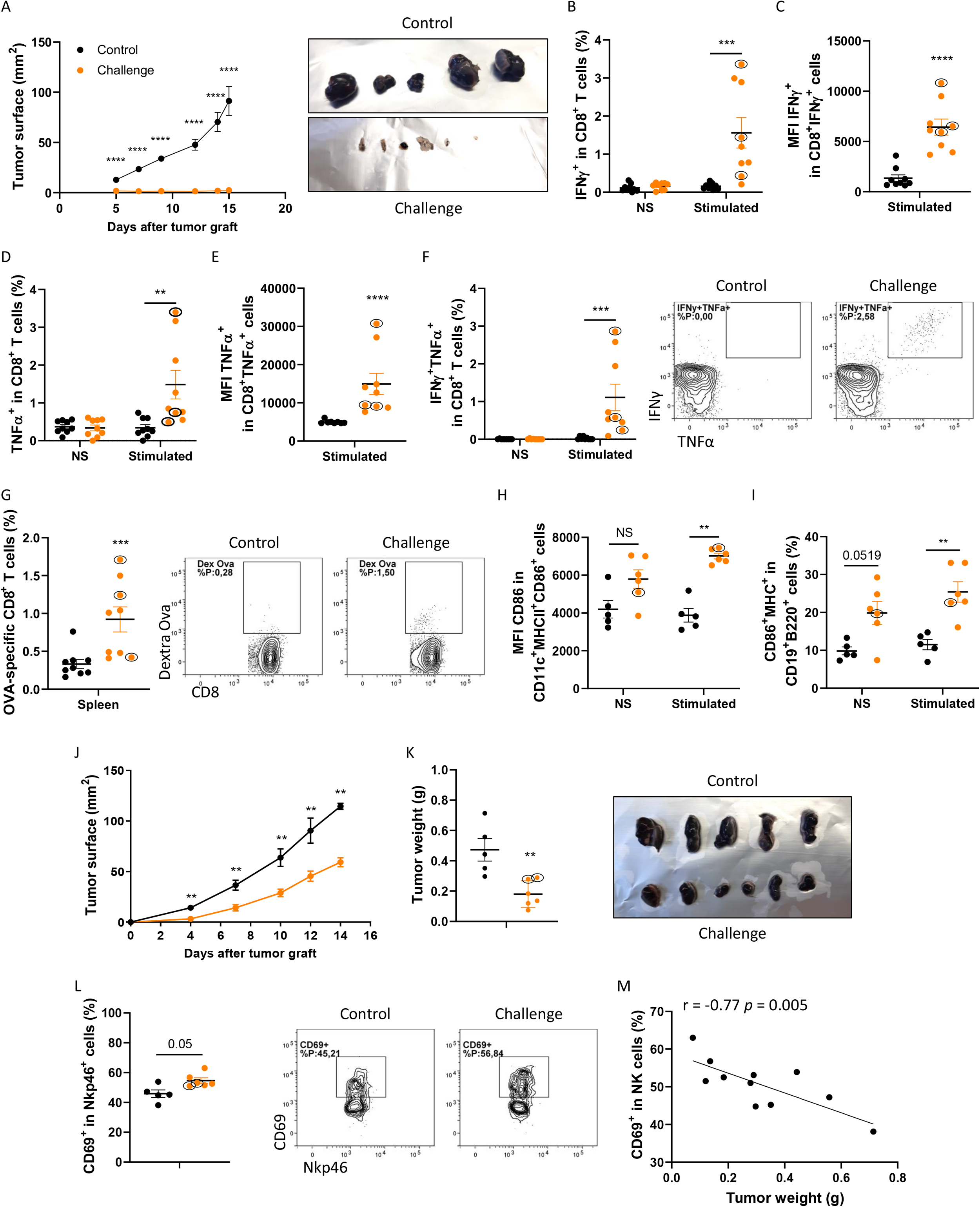
Protection from tumor re-challenge in survivor mice treated with LNP-Ova mRNA +/- anti-PD-1. **(A)** Tumor growth in LNP-Ova mRNA- (orange dot surrounded by a black circle in the following graphs) or LNP-Ova mRNA + anti-PD-1-(orange dot in the following graphs) treated surviving mice challenged with a second injection of B16OVA cancer cells 103 days following the first B16OVA tumor cell injection. Control mice were similar age than challenged mice and were injected with the same number of B16OVA tumor cells (2 × 10^5^). **(B-I)** Frequency of CD8^+^ T cells expressing IFNγ (B,C), TNFα (D,E), coexpressing IFNγ and TNFα (F); frequency of OVA-specific CD8^+^ T cell (G); expression level of CD86 (MFI) in CD86^+^ DCs (H) and frequency of B cells expressing CD86 and MHC-II (I) into the spleens from control and challenged B16OVA tumor-bearing mice following OVA peptide stimulation. **(J**,**K)** Tumor growth (J) and weight (K) in LNP-Ova mRNA- or LNP-Ova mRNA + anti-PD-1-treated surviving mice challenged with an injection of B16F10 cancer cells 113 days following the first B16OVA tumor cell injection. **(L**,**M)** Intratumoral frequency of NK cells expressing CD69 from control and challenged B16F10 tumor-bearing mice (L) as well as correlation between B16F10 intratumoral frequency of CD69^+^ NK cells and tumor weight at day 14 (M). Mean ± SEM of n = 9 mice per group (A-I) or n = 5-6 mice per group (J-M). P values (\**p* < 0.05, \*\**p* < 0.01, \*\*\**p* < 0.001, \*\*\*\**p* < 0.0001) determined by Mann-Whitney test (A-L) or Pearson r correlation test (M).

## Discussion

Therapeutic cancer vaccine is a potential strategy to convert poorly immunologic tumors resistant to anti-PD-1 into inflamed immune tumoral microenvironments favorable for anti-PD-1 efficacy. Here, we demonstrated that mRNA-based vaccine activates DCs *in vitro* in TLR and ROS-dependent signaling pathways and prevents tumor growth in a CD8^+^ T cell-dependent manner in association with an intratumoral IFNγ and antigen presentation gene signature previously shown to correlate with responsiveness to PD-1 blockade in melanoma patients (8,42). We also found that the blockade of PD-1 in vaccinated mice partially prevents the vaccine-related intratumoral increase of PD-1^+^TIM-3^+^CD8^+^ T cell frequency, improves the early production of GzmB by intratumoral CD4^+^ T cells, the survival and the rate of complete tumor regression. Importantly, we observed resistance to tumor growth in surviving mice reexposed to cancer cells expressing or not OVA illustrating immunologic memory establishment and antigen spreading.

DCs are main players in initiating the LNP-mRNA vaccine-induced antitumoral immune response thanks to their ability to detect pathogen-associated molecular patterns (PAMPs), notably nucleic acids and lipids, which activate TLRs leading to the triggering of pro-inflammatory reactions and the subsequent shaping of T cell differentiation (45,46). We should recall that membrane/endosome-localized TLR4 has not been found involved in the mRNA-based vaccine response consensually (24,25) and the functionality of the endosome-localized TLR8 has been described to be controversial in mice (47). Here, we observed a TLR4- and TRL8-dependent DC activation following exposure to LNP-Ova mRNA. This is probably linked to the cationic lipid composition of the Lipidots^®^ and the ssRNA form of mRNA, respectively (46). It has been reported that, TLR agonists such as R848 (TLR7/8) or LPS (TLR4) can trigger ROS-dependent activation and maturation of DCs (44) and that ROS production in DCs plays a crucial role for the induction of CD8^+^ T cell responses following *in vivo* immunization (44) as well as for the proliferation of CD4^+^ T cells following *in vitro* TLR activation (48). Consistent with these studies, we found that DC exposition to LNP-Ova mRNA leads to a NADPH oxidase and mitochondria derivative ROS-dependent activation besides generating OVA-specific CD4^+^ and CD8^+^ T cell activations. Altogether this highlights the potential crosstalk between TLR and ROS signaling pathways in DC response to mRNA-based vaccine leading to the downstream activation of the adaptative immunity.

In this study we used an unmodified mRNA due to its ability to induce signaling by type I IFN which is necessary for the induction of a protective CD8^+^ T cell response (33). Compared with modified mRNA encoding OVA, unmodified mRNA can induce higher frequencies of GzmB^+^/IFNγ^+^/TNFα^+^ polyfunctional OVA peptide-specific CD8^+^ T cells into the spleen and intratumoral activated DCs in association with a robust antitumoral effect as reported in the study of Sittplangkoon C.et al. (33). However, in another study, even if unmodified mRNA leads to higher level of circulating OVA-specific CD8^+^ T cells, better shrinkage of tumor growth is not observed in comparison to a modified mRNA (26). We have shown previously (41) and confirm here that the LNPs we use in this study, which correspond to a second generation of LNP with a lipid core, do not activate DCs solely by themselves unlike LNPs showing intrinsic adjuvant activity. Although modified mRNA improves the translation into protein antigen, its complexation with our LNPs could require the addition of an adjuvant to present a complete vaccinal strategy with optimal antitumoral impact.

Here we report that, besides having *in vivo* prophylactic efficacy in association with the generation of splenic polyfunctional CD8^+^ T cells, LNP-Ova mRNA vaccine displays a CD8^+^ T cell-dependent antitumoral therapeutic efficacy through the robust intratumoral infiltration of OVA-specific polyfunctional CD8^+^ T cells. Our results are consistent with previous preclinical studies showing that mRNA-based vaccines delivering mRNA coding for OVA or tumor antigens (e.g. GP70, TRP-1, TRP-2, MART1, MUC1, E6 or E7, etc.) reduce tumor burden (23–33). Although some of them demonstrate a CD8^+^ T cell infiltration into the tumor (30–33), for the most part, the functionality of CD8^+^ T cells (production/secretion of IFNγ and/or TNFα and/or GzmB) was rather assessed at the splenic or lymph node level (24,25,28–30,32,33) than at the tumor level (23,27). Beyond the implication of CD8^+^ T cells in mRNA vaccine-induced immunity, the intratumoral infiltration of NK cells has been reported with the use of an unmodified mRNA-LNP vaccine in B16OVA tumor-bearing mice (31) as well as in other cancer models submitted to mRNA vaccine (23). In addition to observe a higher GzmB production in NK cells at an early time point (day 9), we also find an increase of their intratumoral frequency, however, their blockade does not impair the vaccinal therapeutic efficacy in our conditions. The early response of NK cells to Ova mRNA vaccine may be not sustained over time to produce a dependent antitumoral effect because of their dysregulation within the TME as recently proposed (49).

Intratumoral cell population immunophenotyping and transcriptional analyses reveal that our mRNA vaccinal strategy promotes a proinflammatory TEM favorable to the transient control of tumor growth while inducing escape mechanisms involving in particular the PD-1/PD-L1 signaling pathway. Blocking PD-1 in addition to the vaccination considerably improves the survival and the rate of complete tumor regression without, at the early time point assessed, deeply impacting the myeloid cell compartment which is mainly affected by the mRNA vaccine. In line with our *in vitro* data and past preclinical works (23,24,33), LNP-Ova mRNA increases the frequency of mature DCs expressing CD86 while inducing a transcriptional type I IFN signature into the tumor. Type I IFN signaling pathway was shown to support the cross-priming of antigen-specific CD8^+^ T cells following DC activation by engagement of TLR such as TLR4 and TLR7/8 (50). The involvement of this mechanism in the antitumor efficacy of LNP-Ova mRNA, although not directly assessed *in vivo*, would be coherent especially in regards to the *in vitro* vaccine-caused TLR-dependent DC activation we observe. Furthermore, LNP-Ova mRNA immunization results in an early and robust reduction in the frequency of immunosuppressive CD206^+^ M2-like tumor-associated macrophages (TAMs) whereas that of inflammatory CD86^+^ M1-like TAMs is barely increased. Although these cells have been rarely explored intratumorally in response to anticancer mRNA vaccines (23,32), a similar phenotype skewing of macrophages has been found into the tumors of mice vaccinated with mRNA coding E7 (23). Mechanistically, this phenotype commitment can be related to the IFNγ and/or TNFα signals through the massive intratumoral infiltration of IFNγ and TNFα producing CD8^+^ T cells following vaccination as already reported in a peptide-vaccine study (33,51). Moreover, the antitumor efficacy of transferred antigen-specific CD8^+^ T cells can be promoted by the migration of Ly6C^high^ monocytes into the tumor (52). Interestingly, we also observe an infiltration of inflammatory monocytes in response to LNP-Ova mRNA vaccine concomitantly with better CD8^+^ T cell activity. However, these cells have a particularly high expression of PD-L1 following vaccination suggesting their potential dual role and involvement in the induction of a later CD8^+^ T cell dysfunction related to the resumption of tumor growth in the majority of animals treated only with the mRNA vaccine. Finally, the mRNA vaccine-induced upregulation of PD-L1 on proinflammatory monocytes, but also on DCs, probably linked to the high intratumoral levels of IFNγ, can be considered as an adjuvant strategy to the PD-1/PD-L1 blockade to both sustain an elevated T cell activation while preventing their mutual inhibition highlighting the interest in therapeutic combinations rather than monotherapy once again.

Although CD8^+^ T cells are undoubtedly crucial players in the anti-tumor response during treatment by immunotherapy in general and therapeutic vaccination in particular, the role of CD4^+^ T cells is increasingly investigated (53). Usually, CD4^+^ T cells have been depicted as helper cells, furthermore, results from preclinical and clinical studies reveal their intrinsic cytotoxic properties as well as their capacity to kill directly cancer cells in a MHC class-II context through notably their production of GzmB (54,55). In a new way compared to previous preclinical studies evaluating combinatory approaches (23,27), we observe at an early stage a positive interactive effect between mRNA vaccine and anti-PD-1 treatment on the intratumoral frequency of Gzmb^+^ CD4^+^ T cells suggesting the potential involvement of these cytotoxic cells in the effective control of the tumor progression in response to the combinatory therapy. These results are in line with the better clinical response to the PD-1/PD-L1 axis blockage associated with the presence of cytotoxic CD4^+^ T into the tumor (56). Concerning the CD8^+^ T cells, although their ability to produce high levels of IFNγ and GzmB is still observed in vaccinated mice over time (day 9 and day 17), the frequency of PD-1^+^ CD8^+^ T cells is already doubled on day 9, reinforcing the interest of adding an anti-PD-1 therapy to maintain the initial antitumor effect of the vaccination-related massive infiltration of active cytotoxic CD8^+^ T cells.

Finally, another key observation is that our vaccinal strategy associated with an anti-PD-1 therapy mediates a strong and sustainable OVA-specific CD8^+^ T cell memory preventing the growth of a new OVA-expressing tumor. The antitumoral immune memory is rarely evaluated in the preclinical studies quoted above, however, in two studies mRNA vaccination targeting CT26-M90 neoepitope or E7 oncoprotein eradicates the initial tumor in 60% or 100% of cases respectively and protects mice from re-challenge. In our experiments, although survival mice treated with the vaccine alone are able to mount a memory immune response, the rate of initial complete tumor regression following vaccination is much lower (25%). These differences may be related to the modifications made or not to the mRNA, the adjuvancity of the LNPs as well as the dose which is largely lower in our study when compared to the two others (3 μg here vs. 40 μg). Nevertheless, in most clinical trials therapeutic mRNA cancer vaccines are more likely to perform well in combination with other immunotherapeutic therapies (22,57,58). We also demonstrate that our vaccine strategy sensitizes an anti-PD-1 resistant tumor model to ICB thus supporting the interest in continuing to test these types of combination in patients.

Importantly, as shown with the B16F10 re-challenge experiment, a secondary immune response against non-OVA derivative antigens (antigen spreading) is fostered following initial therapy-mediated tumor destruction which is a required phenomenon for eliciting a durable immune response and improved outcomes in patients undergoing cancer immunotherapy (59,60). While a higher intratumoral frequency of CD69^+^ CD8^+^ T cells in association with the decrease of B16F10 tumor growth in challenged mice was expected, we rather observe a higher frequency of activated NK cells which correlates negatively with the tumor weight. However, since the CD8^+^ T cell production of cytotoxic molecules (IFNγ, GzmB, etc.) has not been evaluated, the involvement of its improvement at this timing or at an earlier time remains possible. Furthermore, the concept of innate immune memory is less well characterized than adaptive immune memory, nevertheless, diverse subtypes of memory NK cells have been described according to the initial activation trigger (61). While the induction of this memory phenotype has been demonstrated by a combination of cytokines *in vitro* with proven antitumor activity *in vivo* (62) as well as in response to viral infection (63), potential induction by immunotherapy has only been exceptionally investigated. To the best of our knowledge, a single paper demonstrates in a new way the role of memory-like NK cells in preventing melanoma lung metastasis through the release of cytokines induced by a combination of STING-LNPs and CpG-ODNs (64). Our results also support this concept and highlight the potential importance of such cells in the prevention of cancer recurrence following combinatory immunotherapy.

To conclude, we demonstrate the capacity of second generation of LNPs (nanostructured lipid carriers) to deliver mRNA into DCs *in vitro* and activate them through TLR and ROS signaling pathways. This easy-to-use LNP, relying on simple mixture between mRNA and LNP, is an attractive platform for developing mRNA-based vaccines allowing antitumoral immune responses. Our in-depth intratumoral investigations of a cold tumor model reveal some particular cellular mechanisms involved in the mRNA vaccine effectiveness such as the massive infiltration of polyfunctional CD8^+^ T cells and proinflammatory monocytes as well as the early DC activation and protumor M2 reduction. However, a mRNA vaccine-induced PD-1/PD-L1 escape mechanism appears which is effectively targetable with an anti-PD-1 therapy to foster complete tumor rejection, durable immune memory and antigen spreading. Finally, our results highlight the interest in deepening the role of cytotoxic CD4^+^ T lymphocytes and NK cells in future studies to optimize the initial and memory antitumoral immune responses of combinatory therapeutic strategies involving mRNA vaccines and ICBs.

## Authors’ Disclosures

No disclosures were reported.

## Authors’ Contributions

**C. Fournier:** conceptualization, methodology, validation, formal analysis, investigation, writing–original draft, writing–review and editing, visualization, supervision, project administration. **M. Mercey-Rejessac:** investigation, writing–review and editing. **V. Derangère:** validation, investigation, resources, visualization. **A. Al Kadi:** investigation, visualization. **D. Rageot:** investigation. **C. Charrat:** investigation. **A. Leroy:** investigation. **M. Escudé:** resources. **S. Escaich:** resources, **F. Ghiringhelli:** investigation, resources, provided critical feedback. **T. Decaens:** provided relevant comments. **F. P. Navarro:** resources, funding acquisition, writing–review and editing. **E. Jouvin-Marche:** funding acquisition, writing– review and editing. **P.N. Marche:** validation, resources, project administration, funding acquisition.

All authors approved the final version of the manuscript.

## Acknowledgements

This work was supported by Inserm Transfert, la Région Auvergne Rhône Alpes, FINOVI and the French Ministry of Higher Education, research and innovation (LipiVAC, COROL project, funding reference N° 2102992411).

We thank the Microcell core facility of the Institute for Advanced Biosciences (UGA - Inserm U1209 - CNRS 5309), especially Mylène Pezert, for her assistance with equipment. This facility belongs to the IBISA-ISdV platform, member of the national infrastructure France-BioImaging supported by the French National Research Agency (ANR-10-INBS-04).

We thank Marie-Line Cosnier for its help in the project tracking for *CEA, LETI*, Grenoble.

## Tables

**Supplemental Table 1:** BMDC and supplement concentrations during the 10 days of expansion.

|  |  | Day 0 | Day 3 | Day 5 | Day 7 | Day 10 |
| --- | --- | --- | --- | --- | --- | --- |
| *Cell Concentration* |  | 0.6 x 10<sup>6</sup> / mL | 0.5 x 10<sup>6</sup> / mL | 0.5 x 10<sup>6</sup> / mL | 0.5 x 10<sup>6</sup> / mL | According to experiment |
| *Supplement concentration* | IL-6 | 5 ng/ml | 2.5 ng/ml | 2.5 ng/ml | - | - |
|  | FLT-3L | 50 ng/ml | 40 ng/ml | 30 ng/ml | 25 ng/ml | 25 ng/ml |
|  | GM-CSF | 5 ng/ml | 5 ng/ml | 5 ng/ml | 5 ng/ml | 5 ng/ml |

**Supplemental Table 2:**
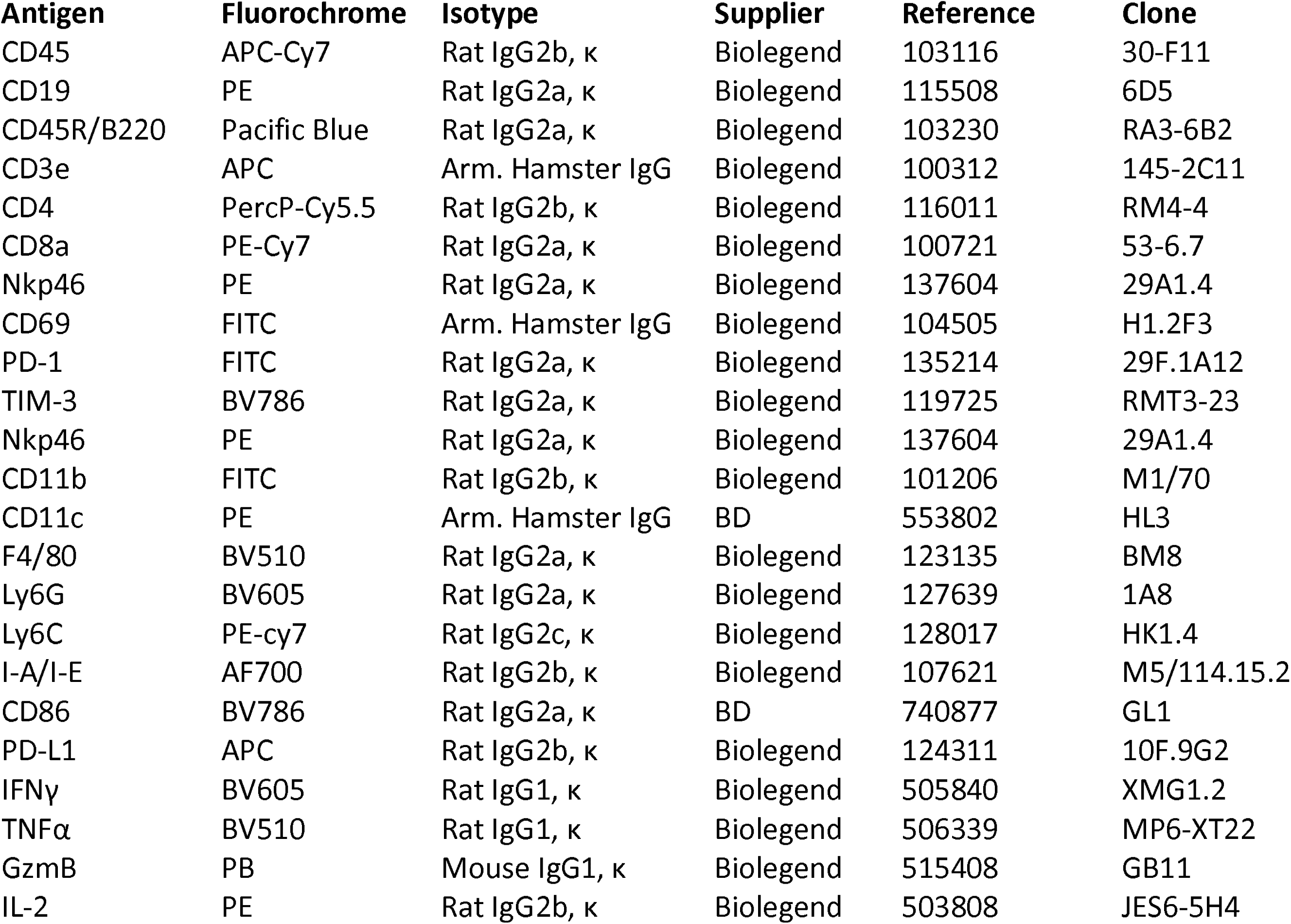
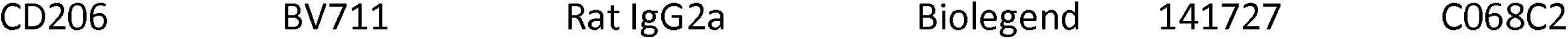
Antibodies used for flow cytometry.

## Figure legends

**Supplemental figure 1: LNP-Ova mRNA mediates BMDC and T cell activation *in vitro* (A**,**B)** Cytotoxicity related to the diverse inhibitors used and the LNP-Ova mRNA treatment was assessed. A positive control was performed with a lysis solution. Mean ± SEM of 2 independent experiments where each dot represents the mean of 2 replicates. **(C-F)** CD8^+^ T cell production and/or secretion of IFNγ (C), TNFα (D,E) and IL-2 (F). (G-I) CD4+ T cell production and/or secretion of IFNγ (G) and TNFα (H,I). Intracellular production was assessed by flow cytometry analysis after intracellular staining and extracellular secretion was evaluated by ELISA on cell supernatants following 48 hours of coculture with BMDCs treated with vehicle, LNP, LNP-cherry mRNA or LNP-Ova mRNA. Mean ± SEM of four independent experiments where each dot represents the mean of 3 replicates. *P* values (\**p* < 0.05, \*\**p* < 0.01, \*\*\**p* < 0.001, \*\*\*\**p* < 0.0001) determined by Kruskal-Wallis test with Dunn’s post hoc testing (C-I).

**Supplemental figure 2: Antitumor effect of LNP-Ova mRNA complex as a preventive vaccine (A)** FMO for determining the positive/negative threshold of IL-2^+^, IFNγ^+^, TNFα^+^, IFNγ^+^TNFα^+^, GzmB^+^, PD-1^+^ CD8^+^ T cell populations. **(B-K)** Following *ex vivo* restimulation with OVA peptides, analysis of cytokine production from splenocyte CD4^+^ T cells (B-D), CD8^+^ (E-G) or CD4^+^ T cells (H-J) from TDLN of B16OVA tumor-bearing mice immunized with LNP-Ova mRNA or LNP-Cherry mRNA by flow cytometry as well as IFNγ secretion in the splenocyte supernatant by ELISA (K). **(L**,**M)** Experimental setting chart (L) and tumor growth in LNP-Ova mRNA or LNP-Egfp mRNA immunized mice injected i.d. with B16OVA melanoma cells 49 days following the prime injection. Mean ± SEM of n = 6 mice per group. *P* values (\**p* < 0.05, \*\**p* < 0.01, \*\*\**p* < 0.001, \*\*\*\**p* < 0.0001) determined by Mann-Whitney test (B-K, M).

**Supplemental figure 3: Antitumor effect of LNP-Ova mRNA complex as a therapeutic vaccine (A**,**B)** Experimental setting charts for engraftment and treatment of B16OVA (A) and E.G7-OVA (B) tumors. **(C**,**D)** Weights of tumor and spleen at day 17 collected from B16OVA-tumor (C) and E.G7-OVA-tumor (D) bearing mice treated with LNP control mRNA or LNP-Ova mRNA. **(E)** Gating strategy for intratumoral CD4^+^ and CD8^+^ T-cell identification using flow cytometry after restimulation with OVA peptides for 4h00. **(F-P)** Number of CD45^+^ cells (F) and CD4^+^ T cells (G) per mg of B16OVA tumor tissue, surface of CD4^+^ T cells analyzed by IHC for each tumor group (H) and representative pictures of PD-1^+^ T cells staining (I), frequency of CD8^+^ T cells expressing IFNγ (J), TNFα (K) and GzmB (L) as well as CD4^+^ T cells expressing CD69 (M), IFNγ (N), TNFα (O) and GzmB (P) from LNP-Ova mRNA or LNP-irrelevant mRNA-treated mice. **(Q-X)** Frequency of CD8^+^ T cells in CD45^+^ cells (Q), frequency of CD8^+^ T cells expressing PD-1 (R), IFNγ and TNFα (S), IFNγ and GzmB (T), frequency of CD4^+^ T cells in CD45^+^ cells (U), frequency of CD4^+^ T cells expressing IFNγ^+^ (V), TNFα^+^ (W) or GzmB^+^ (X). **(Y**,**Z)** Experimental setting chart (Y) and individual tumor growth for each group treated i.p. with LNP-Egfp mRNA or LNP-Ova mRNA as well as with anti-CD8β or its control IgG1 (Z). Mean ± SEM of n = 6 mice per group. P values (\**p* < 0.05, \*\**p* < 0.01, \*\*\**p* < 0.001) determined by Mann-Whitney test (C,D, F-H, J-X)

**Supplemental figure 4: Transcriptional landscape of LNP-Ova mRNA treated B16OVA tumors (A)** Experimental setting chart for B16OVA tumor cell implantation and treatment with LNP-Cherry mRNA or LNP-Ova mRNA before RNA sequencing analysis. **(B)** DEGs statistically up or down regulated between LNP-Cherry mRNA and LNP-Ova mRNA groups following a differential expression analysis performed with DESeq2 software on Dr. Tom (BGI Genomics). **(C)** KEGG pathway relationship network based on the 123 genes related to the “immune system” pathway from the KEGG pathway classification. **(D)** Heatmap of genes related to “antigen processing and presentation” following the selection of the most significative pathways from a KEGG pathway relationship network based on the 123 genes related to the “immune system” pathway. N = 4 mice per group. **(E)** Heatmap of genes corresponding to human gene signature from melanoma patients predicting clinical response to anti-PD-1 blocking antibody (8). Genes are *IDO1, CXCL10, CXCL9, HLA-DRA* (in mice we selected MHC-II genes including *H2-Eb1, H2-Aa, H2-Eb1*), *STAT1* and *IFNG*. N = 4 mice per group.

**Supplemental figure 5: LNP-Ova mRNA vaccine cooperates with anti-PD-1 to suppress B16OVA tumor growth (A)** Flow cytometry gating strategy showing the identification of intratumoral inflammatory monocytes (viable, single CD45^+^ F4/80^-^ CD11c^-^ CD11b^+^ Ly6G^-^ Ly6C^+^) expressing PD-L1 from E.G7-OVA tumors. **(B**,**C)** Frequency of inflammatory monocytes expressing PD-L1 (B) and expression level of PD-L1 (MFI) in PD-L1^+^ inflammatory monocytes (C) from E.G7-OVA tumors treated with LNP-Egfp mRNA or LNP-Ova mRNA at day 17. Mean ± SEM of n = 6 mice per group. P values (\**p* < 0.05) determined by Mann-Whitney test **(D)** Experimental setting chart for B16OVA tumor cell implantation and treatment with LNP-Ova mRNA or LNP-Egfp mRNA as well as with anti-PD-1 or its control IgG2a. **(E)** Individual tumor growth for each group treated i.p. with LNP-Egfp mRNA or LNP-Ova mRNA as well as with anti-PD-1 or its control IgG2a. N = 12 mice per group. **(F)** Flow cytometry gating strategy showing the identification of intratumoral inflammatory monocytes (viable, single CD45^+^ F4/80^-^ CD11c^-^ CD11b^+^ Ly6G^-^ Ly6C^+^) expressing PD-L1, DCs (viable, single CD45^+^ Ly6G^-^ F4/80^-^ CD11c^+^ MHCII^+^) expressing CD86 and macrophages (viable, single CD45^+^ Ly6G^-^ CD11c^-^ CD11b^+^ F4/80^+^) expressing PD-L1, CD86 or CD206 from B16OVA tumors. **(G)** Flow cytometry gating strategy showing the identification of intratumoral CD8^+^ T cells (viable, single CD45^+^ Nkp46^-^ CD3^+^ CD8^+^) coexpressing IFNγ and GzmB and CD4^+^ T cells (viable, single CD45^+^ Nkp46^-^ CD3^+^ CD4^+^) expressing GzmB from B16OVA tumors. **(H-K)** IFNγ serum concentration (H), IFNγ production in intratumoral CD4^+^ T cells (I), intratumoral frequency of NK cells (Nkp46^+^CD3^-^) in CD45^+^ cells (J) and GzmB production in intratumoral NK cells (K) from B16OVA tumor-bearing mice treated i.p. with LNP-Egfp mRNA or LNP-Ova mRNA as well as with anti-PD-1 or its control IgG2a. **(L)** Survival in B16OVA tumor-bearing mice treated i.p. with LNP-Egfp mRNA or LNP-Ova mRNA as well as with anti-Nk1.1 or its control IgG2a. Mean ± SEM of n = 6 mice per group. P values (\**p* < 0.05, \*\**p* < 0.01, \*\*\**p* < 0.001) determined Kruskal-Wallis test with Dunn’s post hoc testing (H) or two-way ANOVA with Tukey’s post hoc testing (I-K) or log-rank (Mantel-Cox) test (L).

**Supplemental figure 6: Protection from tumor re-challenge in survivor mice treated with LNP-Ova mRNA +/- anti-PD-1 (A)** Flow cytometry gating strategy showing the identification of DCs (viable, single CD45^+^ CD19^-^ Ly6G^-^ F4/80^-^ CD11c^+^ MHC-II^+^) expressing CD86 or PD-L1 and B cells (viable, single CD19^+^ B220^+^) coexpressing CD86 and MHC-II into the spleen from control and challenged B16OVA tumor-bearing mice. **(B-G)** Frequencies of DCs (B) and B cells (C) in CD45+ cells; frequency of CD86^+^ cells in DCs (D); representative plots of B cells expressing CD86 and/or and MHC-II (E); expression level of PD-L1 (MFI) in PD-L1^+^ DCs (F) and PD-L1^+^ B cells (G) into the spleen from control and challenged B16OVA tumor-bearing mice. **(H**,**I)** Intratumoral frequency of CD8^+^ T cells in CD45^+^ cells (H) and CD69^+^ CD8^+^ T cells (I) from control and challenged B16F10 tumor-bearing mice. Mean ± SEM of n = 5-6 mice per group. P values (\*\**p* < 0.01) determined by Mann-Whitney test. LNP-Ova mRNA- (orange dot surrounded by a black circle) or LNP-Ova mRNA + anti-PD-1-(orange dot) treated surviving mice.

## Notes

### Competing Interest Statement

The authors have declared no competing interest.

