## Supplemental figures for "Second generation of LNP based mRNA vaccine leads to a T cell–inflamed tumor microenvironment favorable for improving PD-1/PD-L1 blocking therapy and long-term immunity in a cold tumor model"

### Slide 1
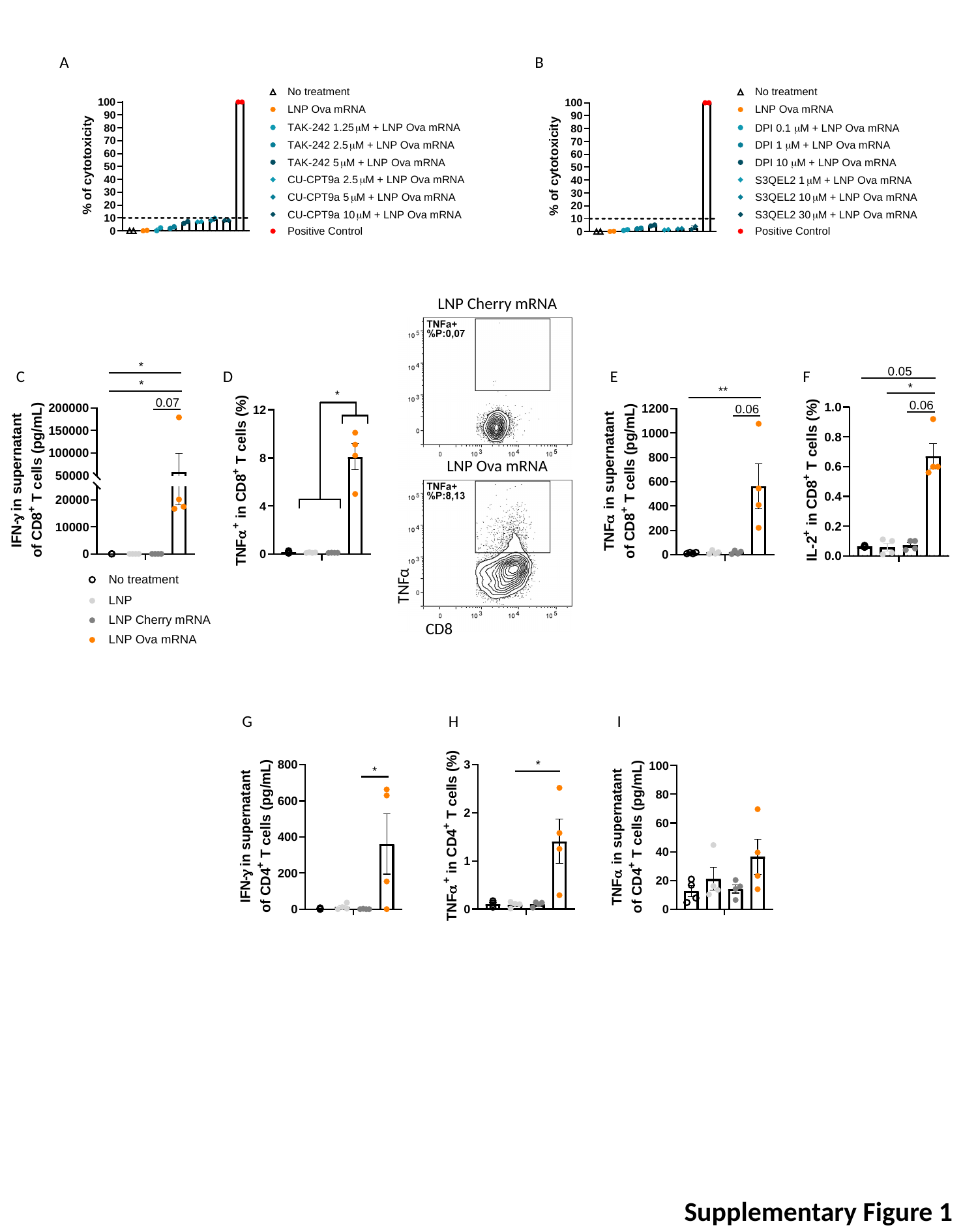

A
B
LNP Cherry mRNA
C
D
E
F
LNP Ova mRNA
TNFα
CD8
G
H
I
Supplementary Figure 1

### Slide 2
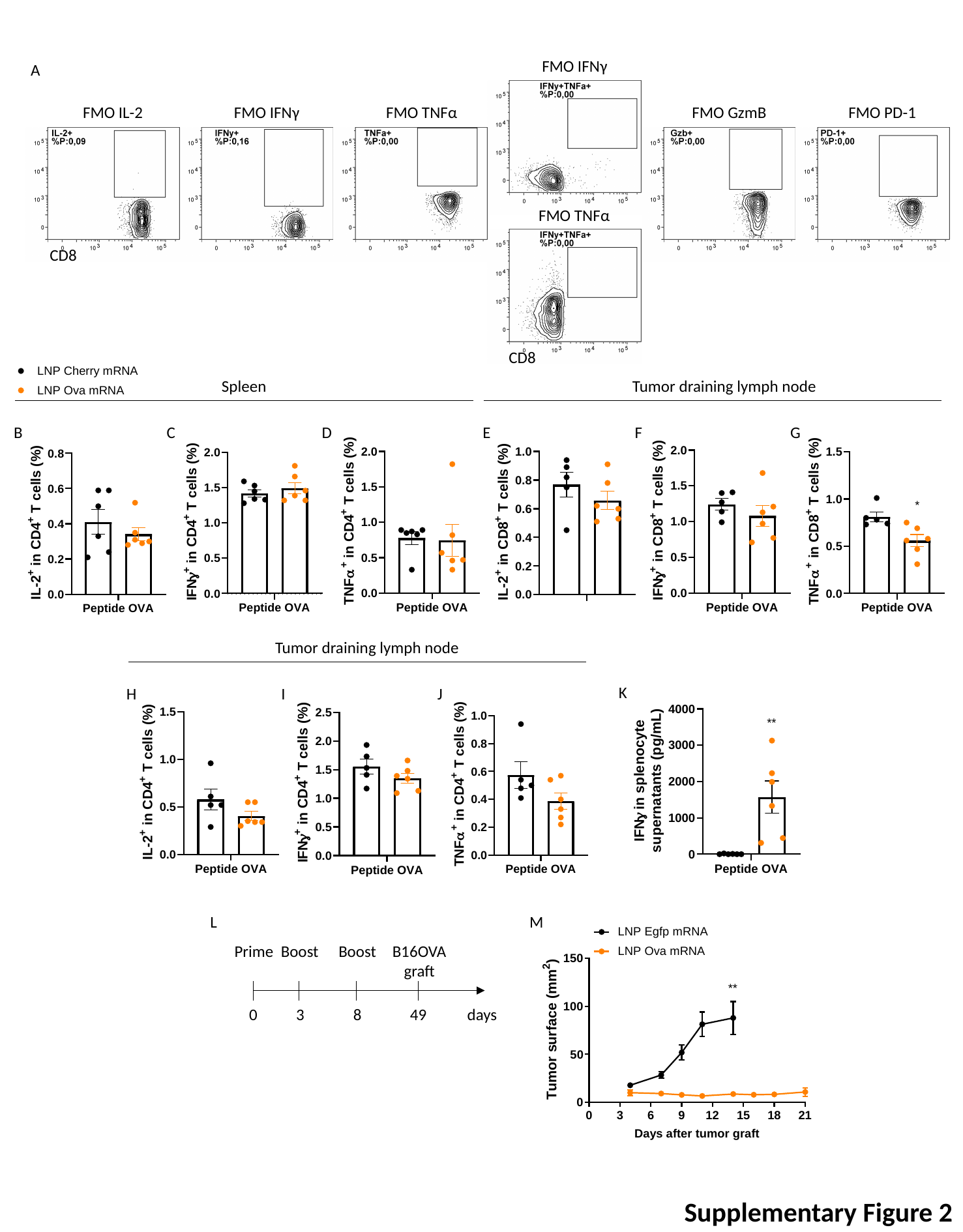

FMO IFNγ
A
FMO IL-2
FMO IFNγ
FMO TNFα
FMO GzmB
FMO PD-1
FMO TNFα
CD8
CD8
Spleen
Tumor draining lymph node
B
C
D
E
F
G
Tumor draining lymph node
K
H
I
J
L
M
Prime
Boost
Boost
B16OVA
graft
49
days
0
3
8
Supplementary Figure 2

### Slide 3
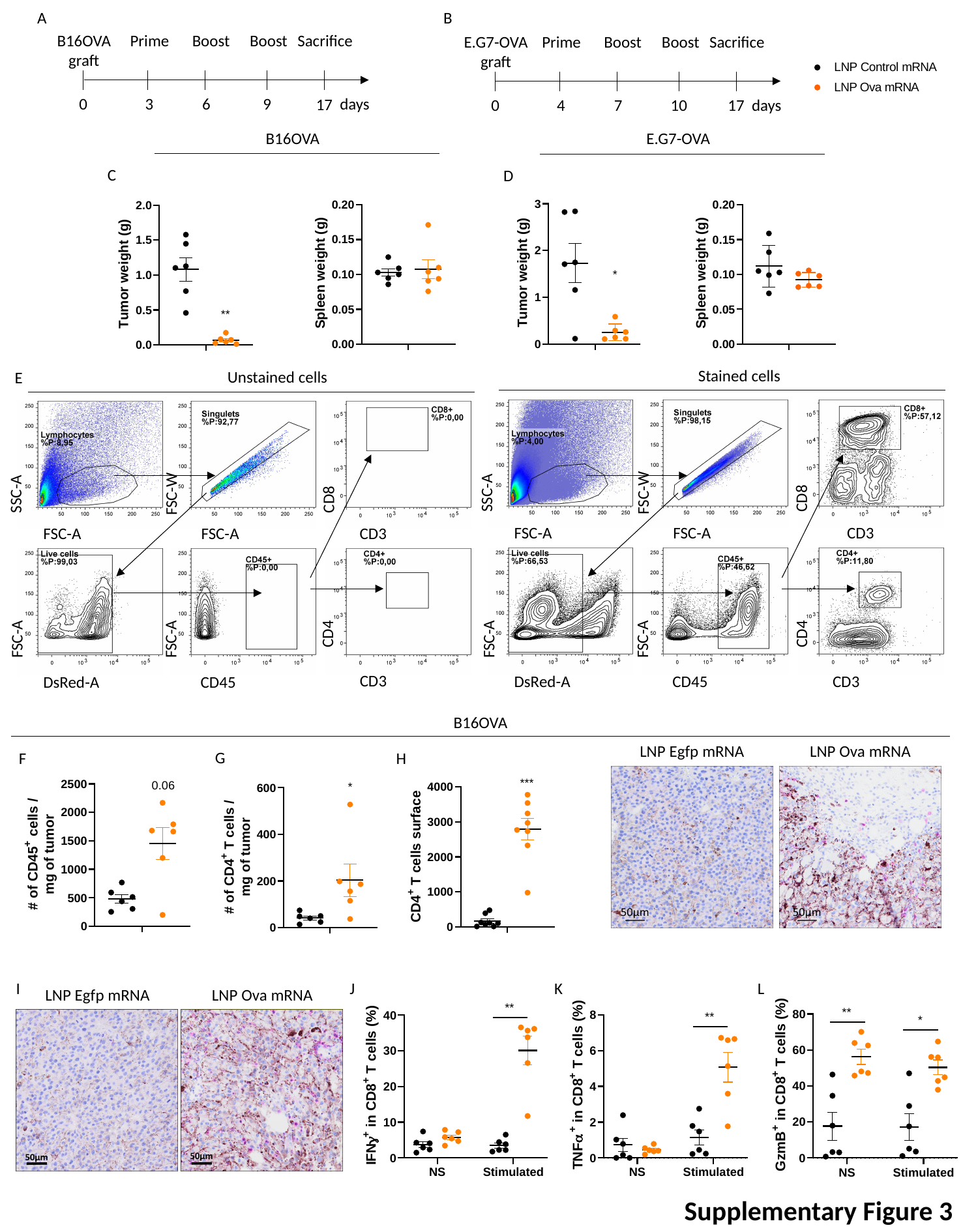

A
B
B16OVA
graft
Prime
Boost
Boost
Sacrifice
9
days
0
3
6
17
E.G7-OVA
graft
Prime
Boost
Boost
Sacrifice
10
days
0
4
7
17
B16OVA
E.G7-OVA
C
D
Stained cells
Unstained cells
E
SSC-A
FSC-W
SSC-A
FSC-W
CD8
CD8
FSC-A
FSC-A
CD3
FSC-A
FSC-A
CD3
CD4
CD4
FSC-A
FSC-A
FSC-A
FSC-A
CD3
CD3
DsRed-A
CD45
DsRed-A
CD45
B16OVA
LNP Egfp mRNA
LNP Ova mRNA
G
F
H
I
J
K
L
LNP Egfp mRNA
LNP Ova mRNA
Supplementary Figure 3

### Slide 4
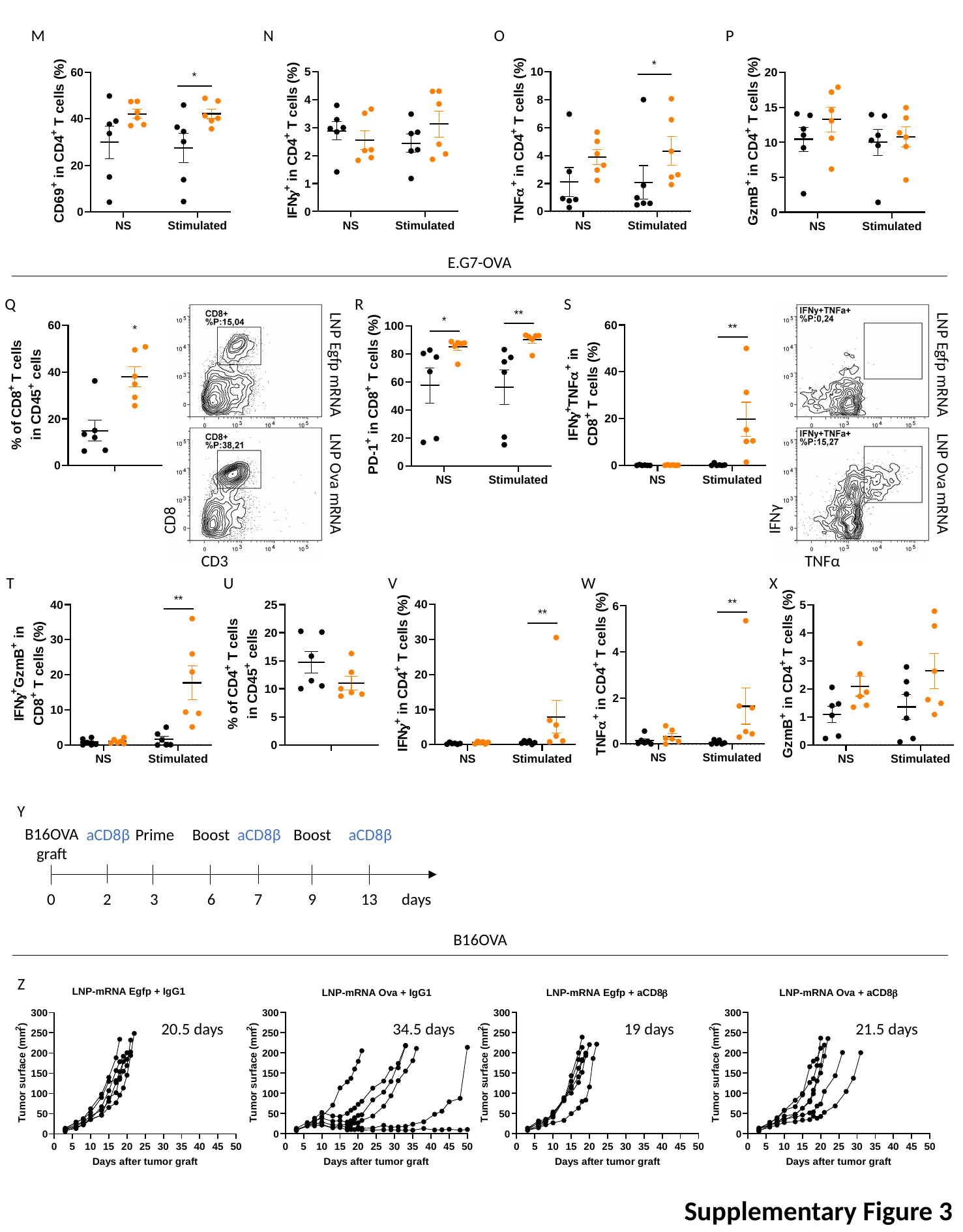

M
N
O
P
E.G7-OVA
Q
R
S
LNP Egfp mRNA
LNP Egfp mRNA
LNP Ova mRNA
LNP Ova mRNA
IFNγ
CD8
N
TNFα
CD3
T
U
V
W
X
Y
B16OVA
graft
aCD8β
Prime
Boost
aCD8β
Boost
aCD8β
0
2
3
6
7
9
13
days
B16OVA
Z
20.5 days
34.5 days
19 days
21.5 days
Supplementary Figure 3

### Slide 5
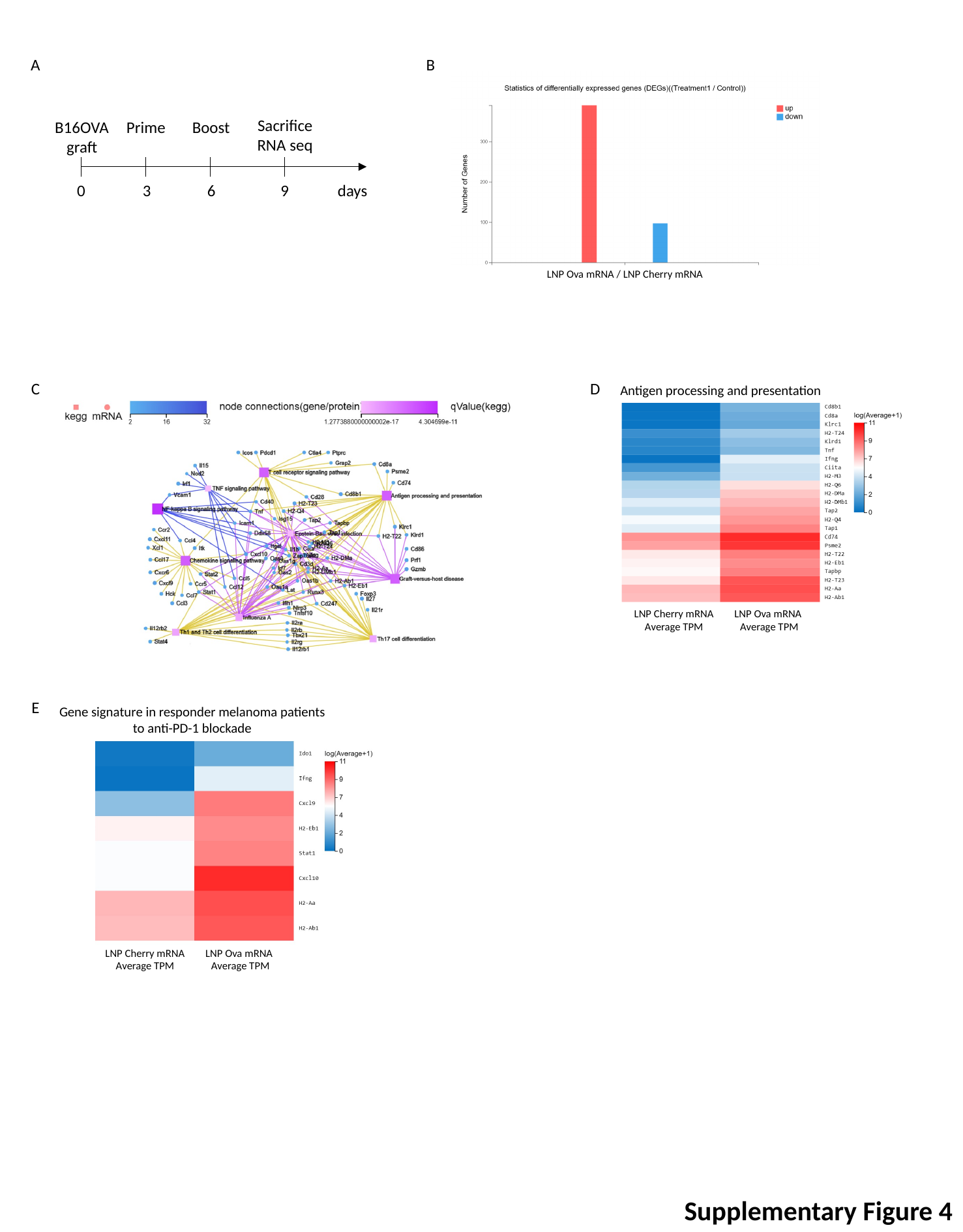

A
B
LNP Ova mRNA / LNP Cherry mRNA
Sacrifice
RNA seq
B16OVA
graft
Prime
Boost
9
days
0
3
6
C
D
Antigen processing and presentation
LNP Cherry mRNA Average TPM
LNP Ova mRNA
Average TPM
E
Gene signature in responder melanoma patients
to anti-PD-1 blockade
LNP Cherry mRNA Average TPM
LNP Ova mRNA
Average TPM
Supplementary Figure 4

### Slide 6
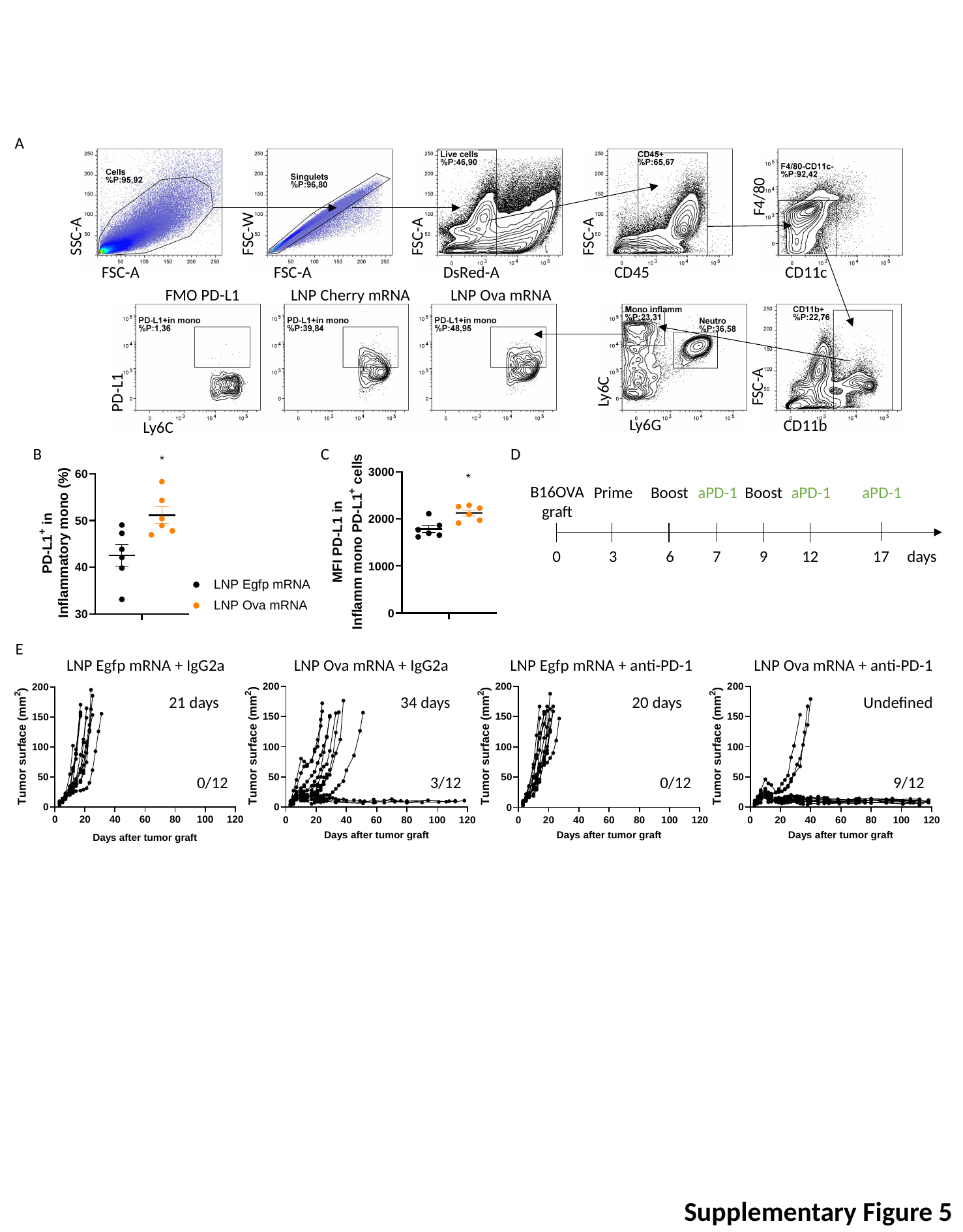

A
F4/80
SSC-A
FSC-W
FSC-A
FSC-A
FSC-A
FSC-A
DsRed-A
CD45
CD11c
LNP Ova mRNA
LNP Cherry mRNA
FMO PD-L1
FSC-A
Ly6C
PD-L1
Ly6G
CD11b
Ly6C
B
C
D
B16OVA
graft
Prime
Boost
aPD-1
Boost
aPD-1
aPD-1
0
3
6
7
9
12
17
days
E
LNP Egfp mRNA + IgG2a
LNP Ova mRNA + IgG2a
LNP Egfp mRNA + anti-PD-1
LNP Ova mRNA + anti-PD-1
21 days
34 days
20 days
Undefined
0/12
3/12
0/12
9/12
Supplementary Figure 5

### Slide 7
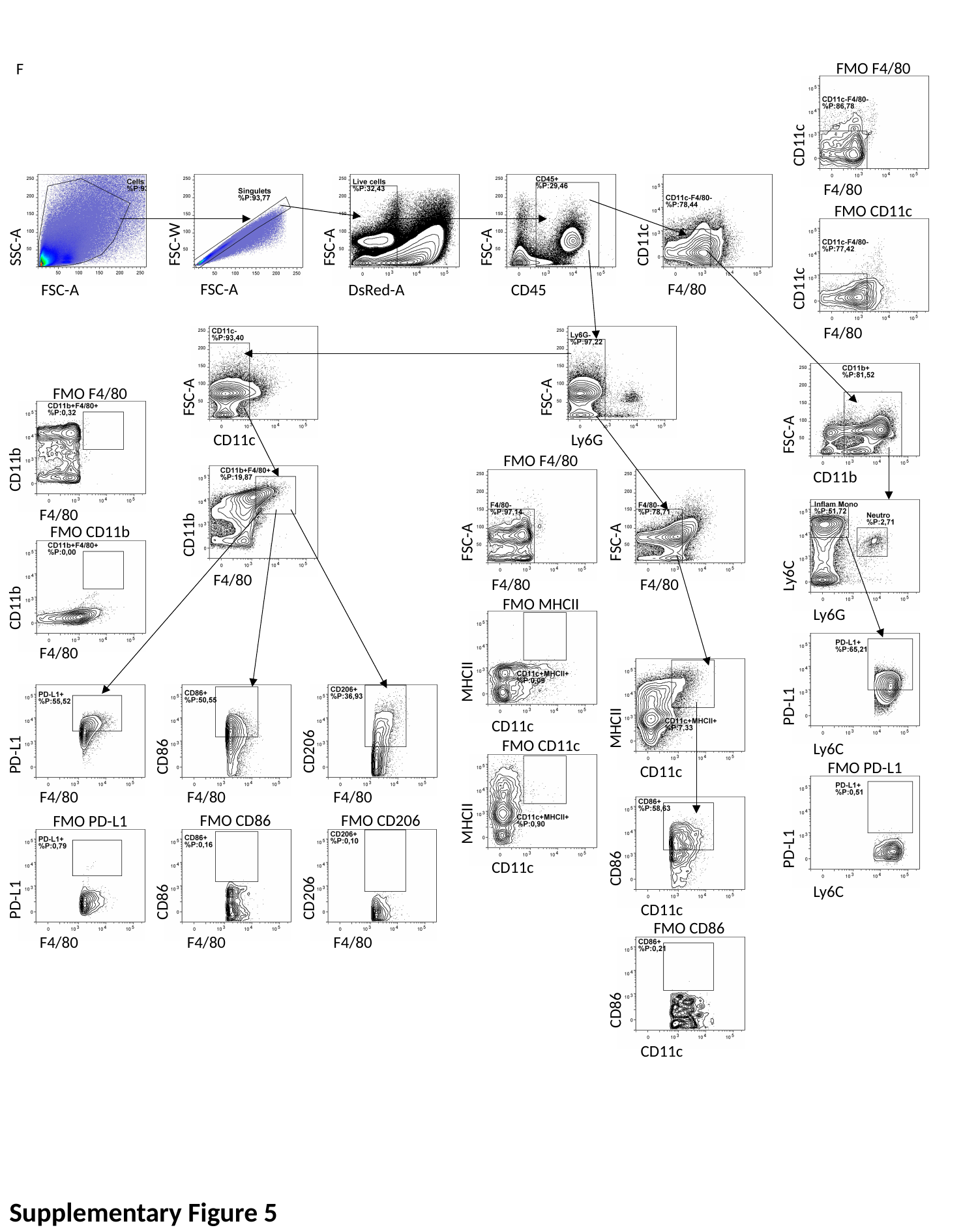

FMO F4/80
F
CD11c
F4/80
FMO CD11c
CD11c
FSC-W
SSC-A
FSC-A
FSC-A
CD11c
F4/80
FSC-A
FSC-A
DsRed-A
CD45
F4/80
FMO F4/80
FSC-A
FSC-A
FSC-A
CD11c
Ly6G
FMO F4/80
CD11b
CD11b
F4/80
CD11b
FMO CD11b
FSC-A
FSC-A
Ly6C
F4/80
F4/80
F4/80
FMO MHCII
CD11b
Ly6G
F4/80
MHCII
PD-L1
MHCII
CD11c
FMO CD11c
CD206
PD-L1
CD86
Ly6C
FMO PD-L1
CD11c
F4/80
F4/80
F4/80
MHCII
FMO CD86
FMO CD206
FMO PD-L1
PD-L1
CD86
CD11c
Ly6C
PD-L1
CD86
CD206
CD11c
FMO CD86
F4/80
F4/80
F4/80
CD86
CD11c
Supplementary Figure 5

### Slide 8
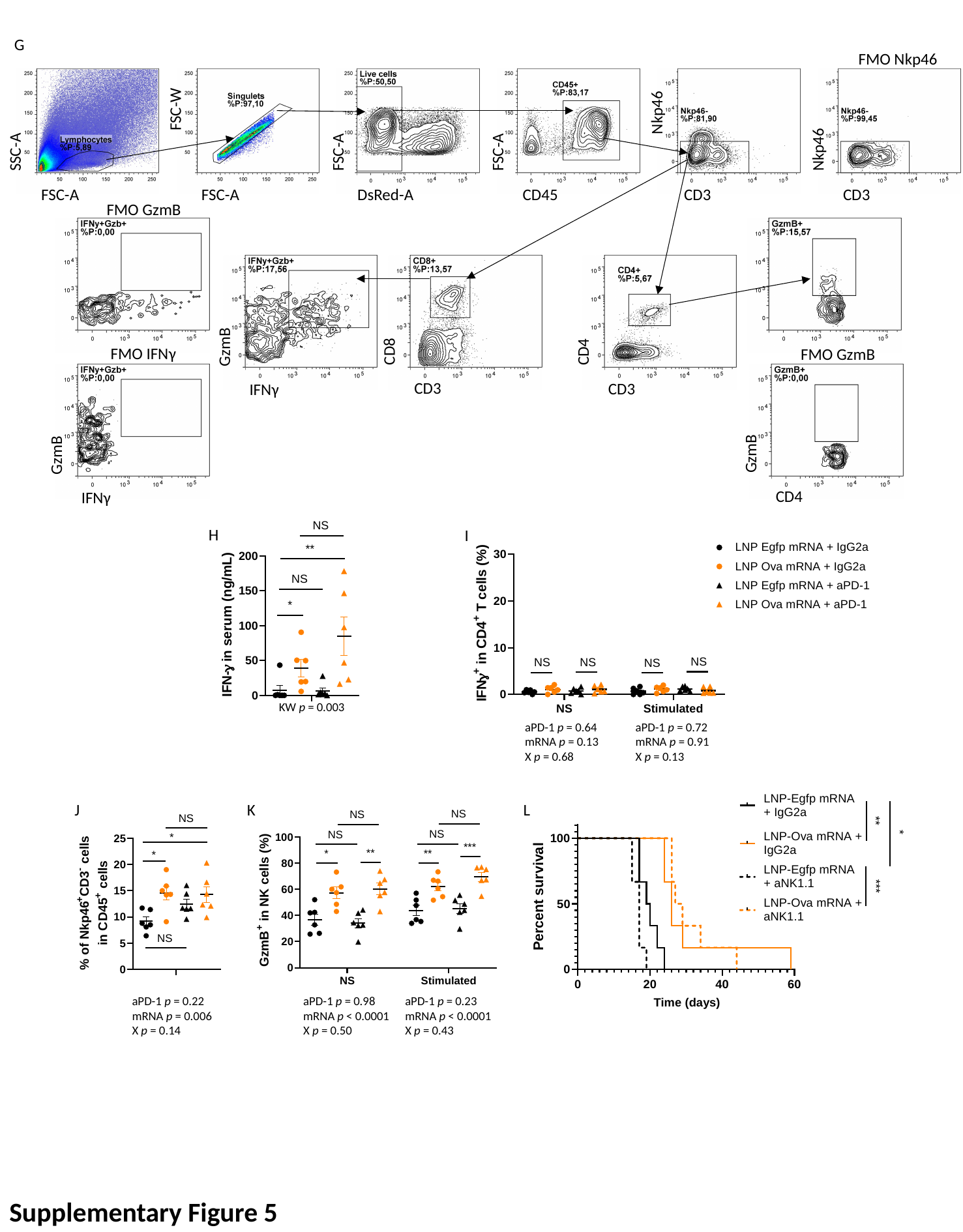

G
FMO Nkp46
Nkp46
FSC-W
Nkp46
SSC-A
FSC-A
FSC-A
FSC-A
FSC-A
DsRed-A
CD45
CD3
CD3
FMO GzmB
CD8
CD4
GzmB
FMO IFNγ
FMO GzmB
CD3
CD3
IFNγ
GzmB
GzmB
CD4
IFNγ
H
I
KW p = 0.003
aPD-1 p = 0.64
mRNA p = 0.13
X p = 0.68
aPD-1 p = 0.72
mRNA p = 0.91
X p = 0.13
J
K
L
aPD-1 p = 0.22
mRNA p = 0.006
X p = 0.14
aPD-1 p = 0.98
mRNA p < 0.0001
X p = 0.50
aPD-1 p = 0.23
mRNA p < 0.0001
X p = 0.43
Supplementary Figure 5

### Slide 9
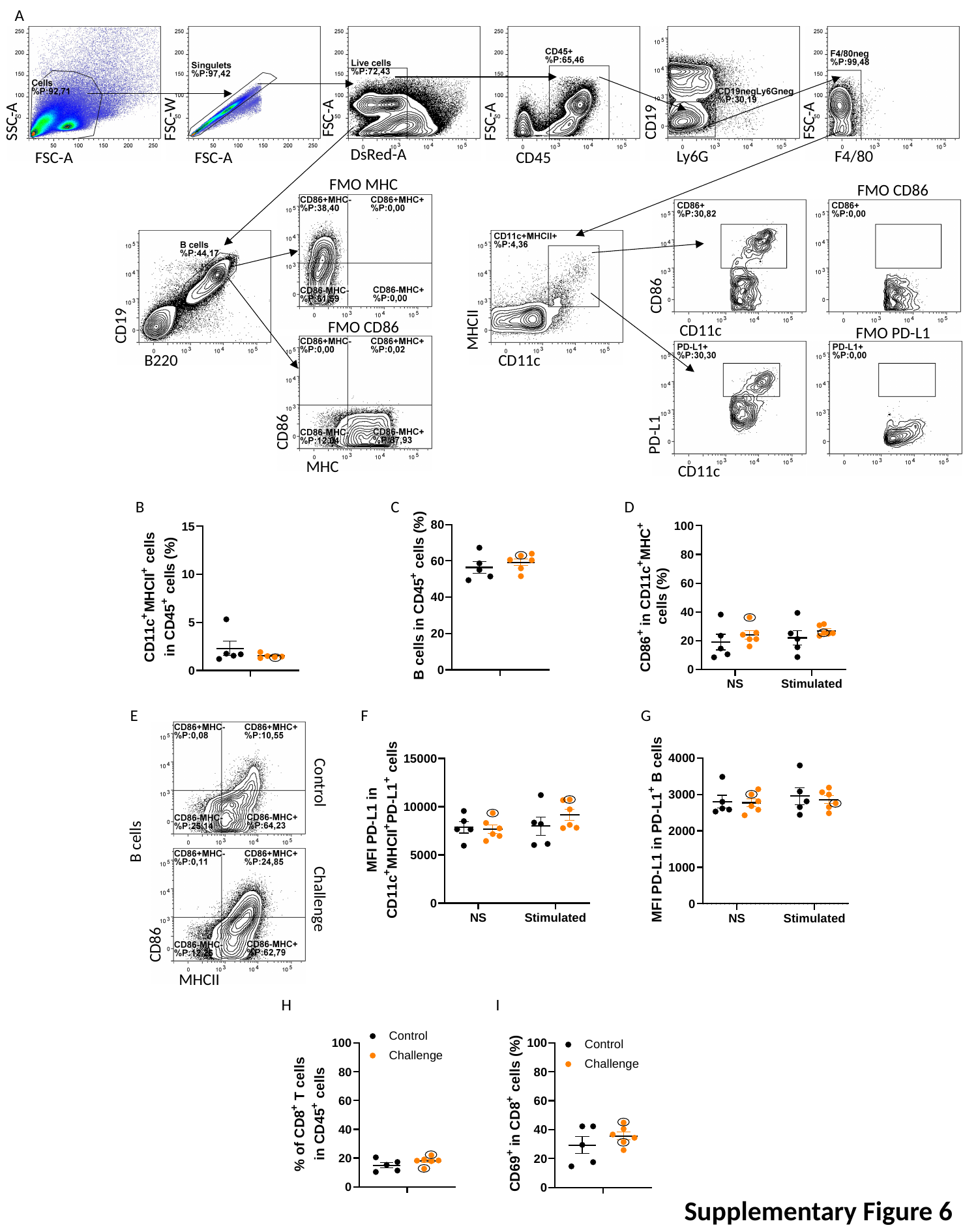

A
SSC-A
FSC-W
FSC-A
CD19
FSC-A
FSC-A
DsRed-A
Ly6G
F4/80
FSC-A
FSC-A
CD45
FMO MHC
FMO CD86
CD86
CD19
MHCII
FMO CD86
CD11c
FMO PD-L1
B220
CD11c
CD86
PD-L1
MHC
CD11c
B
C
D
E
F
G
Control
B cells
Challenge
CD86
MHCII
H
I
Supplementary Figure 6
